# *TWIST1* mediated transcriptional activation of *SPON2* drives colorectal peritoneal metastasis through activation of cancer-associated fibroblast signaling network

**DOI:** 10.1101/2025.04.11.648429

**Authors:** Zhuan Zhou, Alessandro La Ferlita, Manoj H. Palavalli, Xiongfeng Chen, Lauren Tyler, Aslam Ejaz, Joal Beane, Patricio M. Polanco, Huocong Huang, Alex C. Kim

**Affiliations:** Division of Surgical Oncology, Department of Surgery, UT Southwestern Medical Center, Dallas, TX, USA; Division of Medical Oncology, Department of Medicine, The Ohio State University Wexner Medical Center, the James Comprehensive Cancer Center and Solove Research Institute, Ohio State University, Columbus, Ohio, USA; Division of Surgical Oncology, Department of Surgery, The Ohio State University Wexner Medical Center, the James Comprehensive Cancer Center and Solove Research Institute, Ohio State University, Columbus, Ohio, USA; Department of Surgery, Section of General Surgery and Surgical Oncology, University of Illinois-Chicago, Chicago, Illinois, USA

## Abstract

Colorectal cancer (CRC) is the third most commonly diagnosed cancer and the second leading cause of cancer-related mortality in the United States. Peritoneal metastasis (PM), a malignant dissemination within the peritoneal cavity, affects approximately 20% of CRC patients and accounts for 25–35% of stage IV cases. CRC PM is associated with dismal outcomes, with a median overall survival of only 16 months on systemic chemotherapy and an almost 0% five-year survival rate, largely due to frequent treatment resistance and limited therapeutic options.

Despite advances in understanding CRC metastasis, the molecular mechanisms driving CRC PM remain poorly defined. CRC heterogeneity is classified into four Consensus Molecular Subtypes (CMS1-4), with CRC PM tumors predominantly exhibiting the CMS4 signature—characterized by stromal enrichment, high mesenchymal gene expression, and enhanced cellular plasticity—features linked to aggressive disease progression and resistance to standard chemotherapy.

In this study, we identify TWIST1, a basic helix-loop-helix transcription factor, as significantly upregulated in CRC PM. We establish TWIST1-SPON2 as a novel transcriptional axis driving CRC PM tumorigenesis, mediating tumor-stroma interactions between tumor epithelium and cancer-associated fibroblasts (CAFs). Additionally, we identify SPP1, secreted by CAFs, as an upstream regulator of the TWIST1-SPON2 cascade via AKT activation in tumor cells. This newly defined SPP1-TWIST1-SPON2 signaling circuit plays a pivotal role in shaping the tumor microenvironment and promoting CRC PM progression. The findings establish the **SPP1-TWIST1-SPON2** axis as a potential biomarker and a promising therapeutic target in CRC PM.

Keyword: Colorectal cancer, peritoneal metastasis, epithelial-mesenchymal transition, cancer-associated fibroblast

## Introduction

Colorectal cancer (CRC) is the third most common cancer diagnosis and second leading cause of cancer death in the United States [1]. Among its metastatic patterns, peritoneal metastasis (PM) represents a particularly aggressive and challenging form of disease dissemination, affecting approximately 20% of CRC patients and comprising 25– 35% of stage IV cases. PM is associated with the poorest prognosis among metastatic CRC sites, with a median survival of only six months without systemic therapy and 16 months with systemic therapy alone [2]. Despite multimodal treatment approaches, high recurrence rates (50–90%), frequent disease progression, and increasing resistance to therapy contribute to a dismal five-year overall survival rate approaching 0%. With the rising incidence of advanced CRC, particularly early-onset cases (<50 years old), PM represents a growing healthcare burden [1]. Notably, the molecular mechanisms underlying CRC PM remain poorly understood.

A historical paradigm for developing CRC PM envisaged a breach of the colonic wall by T4 (obstructed or perforated) tumors, subsequent exfoliation, and direct peritoneal seeding. However, only 16-19% of T4 patients experience PM, an observation that appears more consistent with the conclusion that PM also arises from T1-3 tumors [3]. These results suggest that CRC PM develops following activation of specific genetic drivers.

The Consensus Molecular Subtype (CMS) classification system categorizes CRC into distinct molecular subgroups, with the CMS4 subtype exhibiting the worst prognosis due to frequent therapeutic resistance and poor relapse-free survival [4]. PMs are highly enriched for the CMS4 signature, which is characterized by upregulation of genes promoting cellular plasticity, metastasis, stromal enrichment, a mesenchymal phenotype, and stemness features [5]. These attributes are strongly associated with high tumor burden and resistance to standard therapies.

Epithelial-to-mesenchymal transition (EMT), a key developmental program aberrantly activated in many cancers, enhances tumor cell plasticity and metastatic potential. EMT is regulated by well-established transcription factors such as TWIST1, SNAI1, SNAI2, and ZEB1. Among these, TWIST1 is a known high-risk factor for CRC metastasis that includes lymph node involvement and tumor budding [6]. Interestingly, TWIST1 activation has been specifically observed in undifferentiated CD44^+^ CRC cells, promoting enhanced tumorigenesis [7]. In this study, we demonstrate the upregulation of TWIST1 in CRC PM and show that TWIST1 knockout (*TWIST1^KO^*) significantly reduces metastatic potential by impairing migration, invasion, and spheroid formation.

Through integrative bioinformatic analyses utilizing chromatin immunoprecipitation sequencing (ChIP-Seq) and RNA sequencing (RNA-Seq), we identified *SPON2* as a direct downstream TWIST1 target and a key regulator of CRC PM tumorigenesis. Additionally, we recently discovered that mesothelial cells are a major source of cancer-associated fibroblasts (CAFs) within the CRC PM tumor microenvironment and secreted phosphoprotein 1 (SPP1) is highly expressed by the mesothelial cell-derived CAFs [8]. into CAFs. Our extensive molecular analyses reveal a novel CRC PM mechanism driven by tumor-fibroblast crosstalk via the SPP1-TWIST1-SPON2 paracrine axis, highlighting a potential therapeutic target for this aggressive disease.

## Materials and Methods

### Biologic sample preparation

Eligible patients underwent informed consent at the Ohio State University Wexner Medical Center under IRB 2019C0139 and 2019C0196. Excess, deidentified tumor specimens were obtained, processed, and entered into the biorepository. Deidentified specimens were utilized to generate protein lysates.

### Cell culture

The cell lines LoVo, CT-26, 3T3, L-WRN, and 293FT were obtained from the American Type Culture Collection (ATCC, Manassas, Virginia, USA). MC38 and MDST8 were procured from Sigma-Aldrich (St. Louis, MO, USA). Cell lines were grown at 37°C in a humidified atmosphere containing 5% CO2. Cultures were maintained in either Dulbecco Modified Eagle Medium (DMEM) (Corning, Corning, NY, USA), F-12 (Gibco, Waltham, MA, USA), or RPMI 1640 (Gibco, Waltham, MA, USA) containing 10% heat-inactivated fetal bovine serum (Sigma-Aldrich, St. Louis, MO, USA), 15 mM HEPES, L-glutamine, and 1% penicillin-streptomycin (Gibco, Waltham, MA, USA). Subculturing of 80%–90% confluent cells was routinely performed using trypsin-EDTA solution (0.25% trypsin and 0.53 mM EDTA).

Pancreatic mesothelial (PanMeso) cells were established and cultured as previously reported [9]. Omental mesothelial (OmenMeso) cells were established as follows. In brief, mesothelium was harvested from the normal omentum of immortomice under a dissecting microscope. The omental mesothelium was seeded onto a tissue culture dish and cultured at 33°C. Expanded cells were collected and named OmenMeso cells. PanMeso and OmenMeso cells were cultured in mesothelial cell media (medium 199 (Gibco/Thermo), 10% FBS, 1% penicillin-streptomycin (Gibco/Thermo), 3.3 nM mouse epidermal growth factor (Biolegend), 400 nM hydrocortisone (MilliporeSigma), 870 nM zinc-free bovine insulin (MilliporeSigma), 20 mM HEPES (Thermo Fisher Scientific)), maintained at 33°C, washed with PBS three times, and cultured in fresh mesothelial cell media at 37°C for three days before any experiments to inactivate immortalization traits. Cells were regularly tested for mycoplasma infection and confirmed to be mycoplasma-free using the e-Myco kit (Boca Scientific).

### shRNA knockdown of TWIST1

Lentiviral short hairpin RNA constructs targeting TWIST1 were obtained from Horizon Discovery (Waterbeach, UK). The target sequences were: AGGAAGAGCCAGACCGGCA, TGTCCGCGTCCCACTAGCA, and GCGGCCAGGTACATCGACT. Viral particles were produced using HEK293T cells with the trans-lentiviral packaging system from Horizon Discovery. Transduction was performed on LoVo cells using 4 µg/mL hexadimethrine bromide (Polybrene). Viral induction was carried out for 8 hours before the media was replaced. Knockdown efficiency was assessed at the mRNA level via qRT-PCR 5 days post-transduction using TaqMan probes (Thermo Fisher Scientific, assay ID# HS04989912_s1) for TWIST1 (Thermo Fisher, Waltham, MA, USA).

### CRISPR Cas9 Knockout of TWIST1 and SPON2

CRISPR-Cas9 knockout of TWIST1 in colorectal cell lines MDST8, MC38, and CT-26 was achieved via electroporation. Guide RNAs were sourced from Integrated DNA Technologies (IDT, Coralville, IA, USA). Human guide RNA sequences for TWIST1 included: TTGCTCAGGCTGTCGTCGGC, GCAAGCGCGGGGGACGCAAG, and CGGGAGTCCGCAGTCTTACG. Mouse guide RNA sequences for TWIST1 were: CACGTTAGCCATGACCCGCT, CGGGAGCCCGCAGTCGTACG, and CGCCGCCCGCGAGATGATGC. Cas9 (Invitrogen, TrueCut) was annealed with guide RNAs and the fluorescent marker ATTO550 (Invitrogen, Waltham, MA, USA). Optimization of electroporation settings was performed using GFP transfection efficiency on the NEON system (Invitrogen, Waltham, MA, USA). Post-optimization, electroporation of MDST8, MC38 and CT-26 cells was conducted. Fluorescently labeled cells were sorted by single-cell isolation and cultured as described. Genomic DNA extraction (Thermo Scientific, Waltham, MA, USA) and PCR amplification of the guide RNA site (forward: CCTCCTCCTCACGTCAGGCCAA and reverse: CTTGCTCAGCTTGTCCGAGGGC) followed by Sanger sequencing were used to evaluate knockout efficiency. Synthego Interference of CRISPR Editing was performed to assess knockout efficiency, with Western blotting verifying protein knockout in MDST8, MC38, and CT-26 cells.

Human and mouse SPON2 sgRNA CRISPR/Cas9 All-in-One Lentivector sets were purchased from Applied Biological Materials Incorporated (Richmond, BC, Canada). Mouse SPON2 target sequences were: KO1 GAAAACGTGAGTCTTGCCCT, KO2 TGGAGCCTATCATGGCCAGG, and KO3 CAAACCGATTCTCCCCCCAG. Human SPON2 target sequences were: KO1 CAGATGGACTCTCCCCCAAG, KO2 GGTGATGCTGTATTTGGCCA, and KO3 GCTGTAGTCGGAGCTATGCG. Viral particles were produced using HEK293T cells with the trans-lentiviral packaging system from Horizon Discovery. Transduction was performed on MDST8 and MC38 cells using 4 µg/mL hexadimethrine bromide (Polybrene®). Western blotting verified protein knockout in MDST8, MC38, and CT-26 cells.

### Plasmids

The mouse Spp1 (NM_009263) promoter mCherry reporter (#MPRM80524-LvPM02; GeneCopoeia) and the mouse Spon2 (NM_133903) promoter GFP reporter (#MPRM64354-LvPF02; GeneCopoeia) were purchased from GeneCopoeia. The GFP-SPON2 expression plasmid (#HG12452-ACG) was obtained from Sino Biological Inc. Plasmids were transfected via Lipofectamine 3000 (Thermo Fisher) to study promoter activity and protein expression.”

#### Western blotting

Proteins were extracted using RIPA buffer (Invitrogen, Waltham, MA, USA), and protein concentrations were determined using a Pierce BCA kit (Thermo Fisher, Waltham, MA, USA). Equal amounts of protein were separated on polyacrylamide gels and transferred to nitrocellulose membranes via the wet tank method. Membranes were blocked for 1 hour using 5% non-fat milk, incubated overnight at 4°C with primary antibodies, washed with TBST, and incubated for 1 hour with horseradish peroxidase (HRP)-linked secondary antibodies. Protein bands were visualized using enhanced chemiluminescent detection reagents (ECL) on a LI-COR Odyssey FC. Primary antibodies were used at a 1:1000 dilution, including TWIST1 (Abcam, 50887), SPON2 (Thermo Fisher PA-106790), SPP1 (InVivoMAb BE0382), p-AKT (Ser473) (Cell Signaling, 3700S), AKT (Cell Signaling, 9272S), p44/42 MAPK (Erk1/2) (Thr202/Tyr204) (Cell Signaling, 4370S), p44/42 MAPK (Erk1/2) (Cell Signaling, 4695S), SNAI1 (Cell Signaling, 3879S), SNAI2 (Cell Signaling, 9585S), ZEB1 (Cell Signaling, 3396S) and β-actin (Cell Signaling, 3700S). Secondary antibodies, anti-mouse IgG (Cell Signaling, 7076S) and anti-rabbit IgG (Cell Signaling, 7074S), were used at a 1:2000 dilution.

### Immunohistochemistry analysis

Formalin-fixed, paraffin-embedded tumor tissue sections (5 μm thick) were tested using antibodies against SPON2 (Thermo Fisher, PA-106790), SPP1 (InVivoMAb, BE0382), CD45 (Cell Signaling, 70257S), and COL1A1 (Cell Signaling Technology, 72026S) according to standard procedures. Briefly, slides were deparaffinized in xylene and rehydrated through a series of decreasing ethanol concentrations in water. Antigen retrieval was performed using antigen retrieval buffer (10 mM Tris-HCl, 1 mM EDTA, 10% glycerol, pH 9.0) in a pressure cooker (Biocare Medical) filled with 500 ml of water. Slides were allowed to cool to room temperature for 30 minutes followed by a PBS rinse. Primary antibodies were applied overnight in a humidified chamber at 4°C. Sections were then washed in buffer for 5 minutes, followed by a 30-minute incubation with ImmPRESS Reagent (VectorLabs, #MP7452 and #MP-7451) and then an additional two 5-minute washes. Then, sections were incubated in DAB working solution (Fisher scientific, 50-823-73) until the desired stain intensity (typically 2–5 minutes) was achieved. Finally, samples were washed twice in buffer for 5 minutes, rinsed in tap water, counterstained with hematoxylin, cleared and mounted. For immunofluorescence staining, after rinsing with PBS, slides were stained with Alexa 488-anti-mouse (Cell Signaling Technology, 4408S) and Alexa 488-anti-rabbit (Cell Signaling Technology, 8889S) secondary antibodies and mounted with Fluoromount-G with DAPI (Thermo Fisher, 00-4958-02). The slides were viewed and photographed using an ECHO Revolve Microscope. Quantification of staining was performed using ImageJ software.

### RNA-Seq

#### Library generation and sequencing

RNA was isolated using the Qiagen RNeasy Mini Kit (ID:74104) (Qiagen, Hilden, Germany). mRNA was purified from total RNA using poly-T oligo -attached magnetic beads. After fragmentation, the first-strand cDNA was synthesized using random hexamer primers, followed by the second-strand cDNA synthesis using dUTP for directional libraries. The library was checked using Qubit and RT-PCR for quantification and a bioanalyzer for size distribution detection. Quantified libraries were pooled and sequenced on the Illumina platform NovaSeq 6000 (Illumina, San Diego, CA, USA).

### Data Analysis

Raw sequencing reads in FASTQ format were quality trimmed, and adapters were removed using Trim Galore (v0.6.6) (https://www.bioinformatics.babraham.ac.uk/projects/trim_galore/). Trimmed reads were then mapped to the human genome (HG38 assembly) using HISAT2 (v. 2.1.0). Afterward, the mapped reads in SAM format were converted into BAM format, sorted for coordinates, and indexed using samtools (v.1.6) [10]. Sorted BAM files were finally used as input for featureCounts (v.2.0.0) in order to count the mapped reads to the gene coordinates reported in the GTF annotation file downloaded from GENCODE (v.43). Raw counts were scaled using the Reads Per Million (RPM) formula to filter out low-expressed genes prior to normalization and differential expression analysis. Precisely, all the genes whose geometric mean of the RPM was less than one across all samples were removed. Afterward, raw counts of retained genes were log2-transformed and differential expression analysis was performed using the Limma R package. Genes with a |Log2FC|>0.58 (|Linear FC|>1.5) and an adjusted p-value <0.05 (Benjamini-Hochberg correction) were considered differentially expressed. All the analyses have been performed in R (v. 4.2.2) using the RStudio (v. 2022.12.0) framework.

### ChIP-Seq

#### Library generation and sequencing

LoVo and MDST8 cells were cultured and fixed with formaldehyde. Cells were lysed with lysis buffer and sonicated. TWIST1 antibody (Abcam ab50887) was incubated with lysed cells and a bacterial positive control (Abcam, Waltham, MA, USA). Antibody complexes were then incubated with Protein G magnetic beads. Samples were washed and eluted using a micro-DNA purification kit, and the purified DNA was used for high-throughput sequencing library construction and protein validation of pulldown.

ChIP DNA was used to generate paired-end libraries using standard Illumina® library protocols. After the library is constructed, use Qubit3.0 for preliminary quantification, dilute the library to 1 ng/μL, and then use Agilent 2100 to detect the length of the inserted sequence of the library. After the length of the inserted sequence meets the expectations, the Bio-Rad CFX 96 fluorescence quantitative PCR instrument is used to accurately quantify the effective concentration of the library (the effective concentration of the library is > 10 nM) to ensure the quality of the library. Qualified libraries were sequenced on the Illumina NextSeq 500 platform, and 150-bp paired-end reads were generated.

### Data analysis

Raw sequencing reads in FASTQ format were quality trimmed, and adapters were removed using Trim Galore (https://www.bioinformatics.babraham.ac.uk/projects/trim_galore/). Trimmed reads in FASTQ format were then aligned to the human genome using Bowtie 2 (v.2.4.5) and, following this, they were converted to the BAM format, sorted for coordinates, and indexed using samtools (v.1.6) [10]. Sorted BAM files were finally used as input for MACS2 (v.2.2.7.1) to perform peak calling. Only peaks with a p-value < 0.05 were used for further analysis. Significant peaks were annotated based on their location relative to the TSS of coding-protein genes by CHIPpeakAnno (v.3.30.1) using the genomic coordinates retrieved from ENSEMBL (internal to CHIPpeakAnno)[11]. The visualization of the peaks around the TSS and other genomic features was performed using the ChIPseeker R package (v.1.32.1) [12].

#### Wound Healing Assay

For the wound healing assay, MDST8 or MC38 cells were plated onto 6-well tissue culture plates coated with 50 μg/ml Matrigel (BD Biosciences, San Jose, CA) with or without 100 ng/ml human SPON2 (R&D Systems, 3266-RS-025/C), 100 ng/ml mouse SPON2 (R&D Systems, 6946-RS-025), 100 ng/ml SPP1 protein (R&D Systems, 441-OP-050 or 11437-OP-050), 1 μg/ml SPON2 antibody (R&D Systems, MAB32661 or MAB3266), or 1 μg/ml SPP1 monoclonal antibody (InVivoMAb, BE0382) with vehicle or isotype IgG controls. Once the cell monolayer reached confluency, a 200 μl pipette tip was used to create scratch wounds. The cells were washed with PBS and then cultured in medium containing 10% FBS in a tissue culture incubator. At designated time points post-scratch, plates were washed with PBS, and the wound width was photographed using an ECHO Revolve Microscope at 5x magnification and measured using ECHO Revolve Microscope software.

#### Haptotactic Migration and Matrigel Chemoinvasion Assays

For haptotactic migration and Matrigel chemoinvasion assays, transwells were pre-coated with 100 ng/ml recombinant mouse/human SPON2 or SPP1 protein or 1 μg/ml SPON2 or SPP1 monoclonal antibody, with or without 150 μg/cm² Matrigel. A total of 5 × 10⁴ MDST8 or MC38 cells were added to the transwells and allowed to migrate towards medium containing 10% FBS. After 24 or 48 hours of incubation, cells on the top side of the transwells were removed with cotton swabs. The transwell membrane was fixed with 4% paraformaldehyde for 20 minutes, and migrated cells were stained with crystal violet and counted under a microscope at 100× magnification. Alternatively, migration and invasion quantification were conducted via colorimetric analysis. Transwell plates were destained with 7% acetic acid, and absorbance was measured at a wavelength of 590 nm using a Synergy H1 Microplate Reader.

#### Tumor spheroid formation assay

Spheroids were generated by culturing 120,000 cells in a 6-well ultra-low attachment plate with conditioned media (advanced DMEM/F-12) derived from L-WRN cells [13]. The media was further supplemented with 1.25 mM N-acetylcysteine (Sigma, cat # A9165), 10 mM nicotinamide (Sigma, Cat # N0636), N21-Max Media Supplement (final concentration 1X) (R&D Systems, cat # AR008), N-2MAX Media Supplement (final concentration 1X) (R&D Systems, cat # AR009), and 10 μM Y-27632 dihydrochloride (Sigma, Cat # Y-27632). When required, 100 ng/ml recombinant mouse/human SPON2 or SPP1 protein or 1 μg/ml SPON2 or SPP1 neutralization antibody with vehicle or isotype IgG controls was added to the culture medium. To propagate the spheroids into subsequent generations, they were filtered through a 40 μm cell strainer and briefly trypsinized. After a brief centrifugation, the cell pellets were resuspended in L-WRN media and replated in an ultra-low attachment plate. Spheroids were photographed using an ECHO Revolve Microscope at 5x magnification, and their diameters were measured using ECHO Revolve Microscope software.

#### Animal Model

All experiments involving animals were performed in accordance with the protocol approved by the University of Texas Southwestern Medical Center. C57BL/6 mice and *Spp1* knockout mice (B6.129S6(Cg)-Spp1tm1Blh/J) were obtained from the Jackson Laboratory (Bar Harbor, ME, USA). Six-week-old C57BL/6 and *Spp1* knockout mice were housed in groups of five per cage. Each mouse received an intraperitoneal injection of 50,000 cells from either the *Twist1* or *Spon2* knockout or control MC38 cell lines. Mice were monitored until they exhibited severe illness or expired naturally. At 28 days post-tumor injection, the control group mice showed signs warranting euthanasia. Mice were sacrificed, and autopsies were conducted. Peritoneal carcinomatosis index (PCI) was independently assessed with a scoring system equivalent to the PCI as used in humans by three researchers (ACK, MHP, and IM) to ensure accurate and unbiased evaluation [14, 15].

#### Statistical Analysis

Statistical analyses were conducted using GraphPad Prism 10. To compare the means of two groups, unpaired Student’s t-tests were utilized. For comparisons among multiple groups, a one-way ANOVA (for one independent variable) or a two-way ANOVA (for two independent variables) with Tukey’s multiple comparisons test was applied to all pairwise combinations. All statistical tests were two-sided, and a P value of <0.05 was considered significant. Mortality rates between groups were compared using log-rank tests, with significance also set at a P value of <0.05.

## Results

We analyzed the previously published large CRC PM dataset, *GSE183202,* to investigate the expression pattern of canonical epithelial-to-mesenchymal transition (EMT) factors (TWIST1, SNAI1, SNAI2, ZEB1) in colorectal cancer peritoneal metastases (CRC PM), [5]. Differential gene expression profiling comparing CRC PM samples with unpaired primary tumors revealed a significant upregulation of TWIST1 (p = 0.031) and SNAI2 (p = 0.025), while SNAI1, and ZEB1 revealed no significant changes in expression (Fig. 1A, S1). Western blot analysis of patient-derived CRC PM specimens further validated the increased TWIST1 protein expression (Fig. 1B).

**Figure 1.**
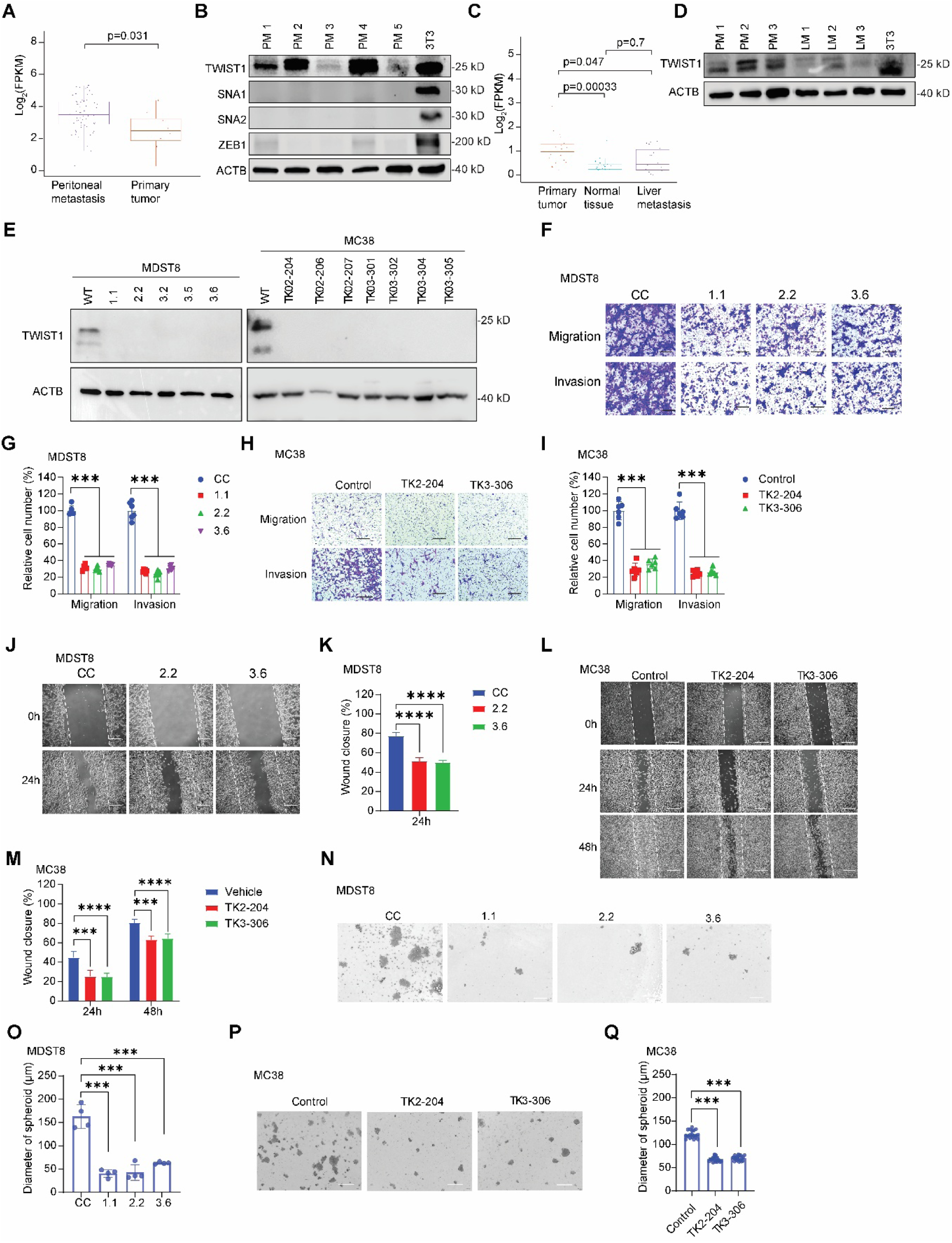
TWIST1 is preferentially expressed in colorectal cancer (CRC) peritoneal metastases (PM) and regulates colon cancer cell migration, invasion, and stemness. (A) Differential expression profiling of CRC PM patient samples reveals upregulation of TWIST1. (B) Western blot analysis of epithelial-mesenchymal transition (EMT) markers TWIST1, SNAI1, SNAI2 and ZEB1 in CRC PM patient samples. (C) Differential expression analysis of TWIST1 in CRC liver metastasis (LM) patient samples shows no significant difference between LM and normal tissue. (D) Western blot analysis of TWIST1 expression in CRC LM and PM samples. (E) Western blot analysis of TWIST1 expression in MDST8 and MC38 TWIST1 single-clone knockout (KO) cells. (F, G) Transwell migration and Matrigel invasion assays were performed to assess haptotactic migration and Matrigel chemoinvasion in MDST8 TWIST1 KO clones. Representative images are shown in (F), with statistical results in (G). Scale bar: 200 µm. (H, I) Transwell migration and Matrigel invasion assays were conducted to evaluate haptotactic migration and Matrigel chemoinvasion in MC38 TWIST1 KO clones. Representative images are shown in (H), with statistical results in (I). Scale bar: 200 µm. (J, K) Wound-healing assays were performed to assess the migration of MDST8 TWIST1 KO clones. Representative images are shown in (J), with statistical results in (K). Scale bar: 200 µm. (L, M) Wound-healing assays were performed to evaluate the migration of MC38 TWIST1 KO clones. Representative images are shown in (L), with statistical results in (M). Scale bar: 200 µm. (N, O) Self-renewal capacity of MDST8 TWIST1 KO clones was assessed. Representative images are shown in (N), with statistical results in (O). Scale bar: 500 µm. (P, Q) Self-renewal capacity of MC38 TWIST1 KO clones was assessed. Representative images are shown in (P), with statistical results in (Q). Scale bar: 500 µm. Results are expressed as mean ± SD. *P < 0.05, **P < 0.01, ***P < 0.001, and ****P < 0.0001.

To assess whether *TWIST1* expression is specific to the peritoneal metastatic site, we examined transcriptomic data from *GSE50760*, which includes colorectal liver metastases (CRLM) and primary tumors [16]. Both transcript and protein levels of TWIST1 were significantly downregulated in CRLM compared to primary tumors (Fig. 1C, D). Collectively, these findings identify TWIST1 as the only canonical EMT factor specifically upregulated in CRC PM, suggesting its potential role in peritoneal metastatic progression.

Next, we performed CRISPR-Cas9-mediated knockout of *TWIST1* (*TWIST1^KO^*) in the human cell line MDST8 and murine cell lines CT26 and MC38, alongside shRNA-mediated *TWIST1* knockdown in the human metastatic CRC cell line LoVo to elucidate the role of TWIST1 in CRC PM tumorigenesis, (Fig. 1E, S2). Analysis of *TWIST1^KO^* monoclonal colonies demonstrated a significant decrease in migration, invasion, and wound closure capacities (Fig. 1F-M, S2), consistent with the established role of TWIST1 in promoting cellular motility and invasive potential essential for the metastatic cascade. Prior studies have shown that TWIST1 regulates the undifferentiated state in primary CD44^+^ CRC cells [7]. As such, we challenged *TWIST1^KO^* cells in a three-dimensional (3D) suspension culture to evaluate their self-renewal capacity via sphere formation. Notably, *TWIST1^KO^* cells exhibited a significant deficit in sphere formation capacity, with decreased spheroid diameter in both MDST8 and MC38 cell lines (Fig. 1M-Q). Collectively, these findings indicate that *TWIST1^KO^*leads to substantial *in vitro* deficiencies in metastasis, characterized by reductions in invasion and self-renewal, which are critical for distant organ metastasis.

To identify the dysregulated downstream target genes responsible for this phenotype, we performed RNA sequencing (RNA-seq) on TWIST1^KO^ and shRNA-TWIST1 knockdown cells (Fig. S3). Concurrently, we conducted chromatin immunoprecipitation sequencing (ChIP-Seq) against TWIST1 in wildtype cells to identify direct downstream target genes (Fig. 2A, S3). Through complex bioinformatics analysis that integrated RNA-seq and ChIP-Seq results (Fig. 2B, S3), we identified 27 dysregulated TWIST1 direct target genes. Notably, the following genes were significantly upregulated in TWIST1^KO^ and shRNA-TWIST1 knockdown cells: *KCNAB2, MIR3142HG, PIK3IP1, PODXL2, SLAIN1, PCDHGC3, MGMT, PDCD4, ACTR3C, TNFRSF21, TNFAIP3, SLC12A2,* and *PPM1N*. Conversely, the significantly downregulated genes in TWIST1^KO^ and shRNA-TWIST1 knockdown cells included *F3, CDC20, SYNE1, SLC7A5, CENPM, ASF1B, SFXN2, DBF4B, SLC7A1, RFX2, LPAR1, FOXL1, TNS1,* and *SPON2* (Fig. 2B, S3).

**Figure 2.**
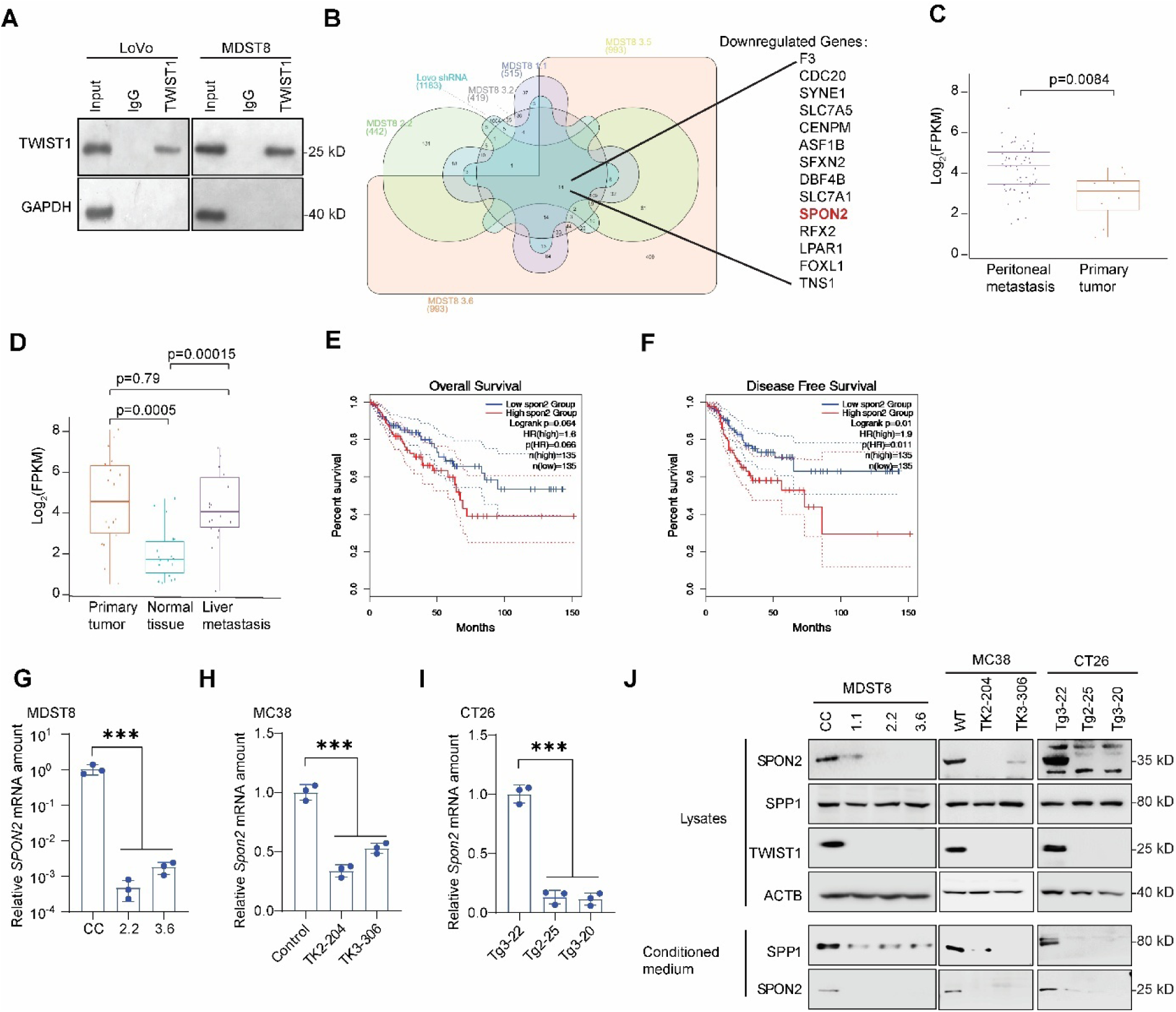
TWIST1 regulates SPON2 expression. (A) Western blot analysis of chromatin immunoprecipitation (ChIP) pull-down experiments for TWIST1 in LoVo and MDST8 cells. (B) Integration of ChIP-seq and RNA-seq data from two metastatic cell lines with biological replicates. Venn diagrams show genes with promoters bound by TWIST1 (ChIP-seq) that are downregulated in TWIST1-deficient LoVo and MDST8 cells. SPON2 is highlighted in red. (C) Differential expression analysis of SPON2 in CRC PM patient samples reveals upregulation. (D) Differential expression analysis of SPON2 in CRC liver metastasis (LM) patient samples. (E) Kaplan-Meier survival analysis of overall survival based on SPON2 expression levels (high vs. low) in The Cancer Genome Atlas (TCGA) patient samples. (F) Kaplan-Meier survival analysis of disease-free survival based on SPON2 expression levels (high vs. low) in TCGA patient samples. (G–I) SPON2 mRNA expression in TWIST1 knockout cells, including MDST8 (G), CT26 (H), and MC38 (I). (J) Western blot analysis of SPON2 and SPP1 expression in whole-cell lysates and conditioned medium from TWIST1 knockout MDST8, MC38, and CT26 cells. Results are expressed as mean ± SD. *P < 0.05, **P < 0.01, ***P < 0.001, and ****P < 0.0001.

We further assessed these 27 target genes in CRC PM relative to primary tumors to identify clinically relevant, dysregulated direct downstream TWIST1 target genes. *SPON2* was identified as a single downstream target gene significantly upregulated in CRC PM compared to primary tumors (P = 0.0084) (Fig. 2C). SPON2 is a secreted protein with a limited understanding of its biological function in development and tumorigenesis [17]. Although *SPON2* expression has been correlated with CRC, its role in metastasis remains relatively unclear. Next, the examination of *SPON2* expression in colorectal liver metastases (CRLM) compared to primary tumors did not reveal preferential expression (Fig. 2D). While there was a trend towards worse overall survival in patients with high *SPON2* levels according to TCGA analysis using *GEPIA 2.0* (https://gepia2.cancer-pku.cn/), a significant decrease in disease-free survival was observed in patients with elevated *SPON2* expression (Fig. 2E-F). Moreover, *SPON2* levels significantly correlated with a higher hazard ratio in advanced stages of CRC in analysis using *TIMER 2.0* (http://timer.cistrome.org) (Fig. S4). Importantly, analyses of human and murine CRC cell lines with *TWIST1^KO^* demonstrated a significant reduction in both mRNA and protein expression of SPON2 (both in lysate and secreted forms), further implicating *SPON2* as a crucial direct target of TWIST1 (Fig. 2G-J). Given the specificity of SPON2 expression in CRC PM, these data suggest the importance of this secreted factor in CRC PM tumorigenesis and patient outcomes.

With a goal of better understanding the biologic implication of SPON2, we performed CRISPR-Cas9 knockout of *SPON2* in both MDST8 and MC38 metastatic CRC cell lines (Fig. 3A). Consistent with the *TWIST1^KO^* phenotype, *SPON2* knockout (*SPON2^KO^*) cells exhibited significant deficiencies in migration, invasion, wound healing, and sphere formation (Fig. 1F-M, 3B-L). We also conducted neutralization of SPON2 protein with monoclonal antibodies, which resulted in significant and analogous reductions in migration, invasion, wound healing, and sphere formation, highlighting SPON2 as a potential therapeutic target (Fig. 3M-P, S5A-F). Conversely, treatment of MDST8 or MC38 cells with recombinant human SPON2 or murine Spon2, respectively, not only accelerated wound closure, invasion, and sphere formation in wildtype cells but also rescued the *TWIST1^KO^*and *SPON2^KO^* phenotypes (Fig. 3M-P, S5A-H). Moreover, overexpression of GFP-tagged SPON2 in *TWIST1^KO^* cells rescued the *TWIST1^KO^* phenotype, further solidifying the critical role of *SPON2* in mediating these cellular processes downstream of TWIST1 (Fig. 3Q-S, S5I-L).

**Figure 3.**
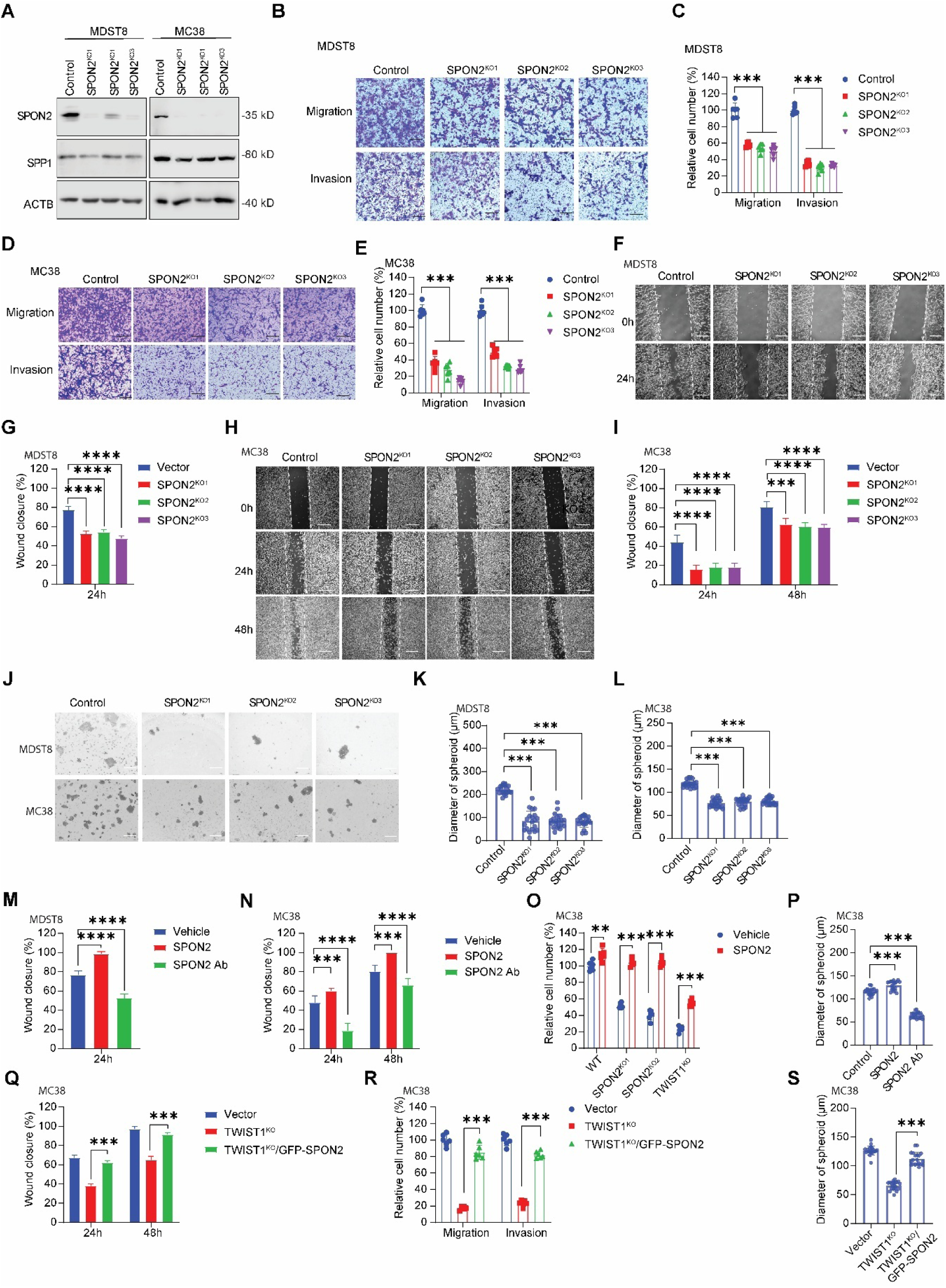
The TWIST1-SPON2 cascade regulates colon cancer cell migration, invasion, and stemness. (A) Western blot analysis of SPON2 expression in MDST8 and MC38 SPON2 knockout (KO) cells. (B, C) Transwell migration and Matrigel invasion assays were performed to assess haptotactic migration and Matrigel chemoinvasion in MDST8 SPON2 KO clones. Representative images are shown in (B), and statistical results are presented in (C). Scale bar: 200 µm. (D, E) Transwell migration and Matrigel invasion assays were conducted to evaluate haptotactic migration and Matrigel chemoinvasion in MC38 SPON2 KO clones. Representative images are shown in (D), with statistical results in (E). Scale bar: 200 µm. (F, G) Wound-healing assay to assess the migration of MDST8 cells plated on Matrigel-coated plates. Representative images are shown in (F), with statistical results in (G). Scale bar: 200 µm. (H, I) Wound-healing assay to assess the migration of MC38 cells plated on Matrigel-coated plates. Representative images are shown in (H), with statistical results in (I). Scale bar: 200 µm. (J–L) Self-renewal capacity of MDST8 and MC38 SPON2 KO clones. Representative images are shown in (J), with statistical results presented for MDST8 in (K) and MC38 in (L). Scale bar: 500 µm. (M, N) Wound-healing assay to assess the migration of MDST8 (M) and MC38 (N) cells plated on Matrigel-coated plates with or without 100 ng/ml SPON2 protein or 1 µg/ml SPON2 monoclonal antibody. (O) Transwell Matrigel invasion assays were conducted to evaluate the effect of SPON2 protein on the chemoinvasion of MC38 TWIST1 or SPON2 KO cells. The Transwell was coated with 1 mg/ml Matrigel with or without 100 ng/ml SPON2 protein and incubated for 48 hours in the presence of 10% FBS. (P) Assessment of self-renewal capacity in MC38 cells cultured with 100 ng/ml SPON2 protein or 1 µg/ml SPON2 monoclonal antibody. (Q) Wound-healing assay to assess the migration of MC38 TWIST1 KO cells with GFP-SPON2 overexpression. MC38 cells were plated on Matrigel-coated plates. (R) Transwell migration and Matrigel invasion assays to assess haptotactic migration and Matrigel chemoinvasion in MC38 TWIST1 KO cells overexpressing GFP-SPON2. (S) Self-renewal capacity of MC38 TWIST1 KO cells overexpressing GFP-SPON2. Results are expressed as mean ± SD. *P < 0.05, **P < 0.01, ***P < 0.001, and ****P < 0.0001.

*TWIST1* expression, specifically in CRC, is an important regulator of tumorigenesis and metastasis through transcriptional regulation of dedifferentiation of the CD44^+^ CRC population [7]. One of the major ligands for CD44 is secreted phosphoprotein 1 (SPP1), which is secreted by cancer-associated fibroblasts (CAFs) in CRC PM [8]. Therefore, we examined whether SPP1 stimulation in tumor cells could activate downstream TWIST1 and subsequent transcription of *SPON2*. Stimulation of MC38 and MDST8 cells with SPP1 led to significant increases in *TWIST1* and *SPON2* transcript levels, protein expression, and SPON2 secretion (Fig. 4A, 4B). We also established stable MC38 cells transfected with pEZX-mSpon2-GFP, a vector with GFP under the control of the *mSpon2* promoter. Treatment of these cells with recombinant SPP1 protein resulted in enhanced GFP expression, confirming the regulation of *Spon2* gene transcription by SPP1 in cancer cells (Fig. 4C-D, S6A).

**Figure 4.**
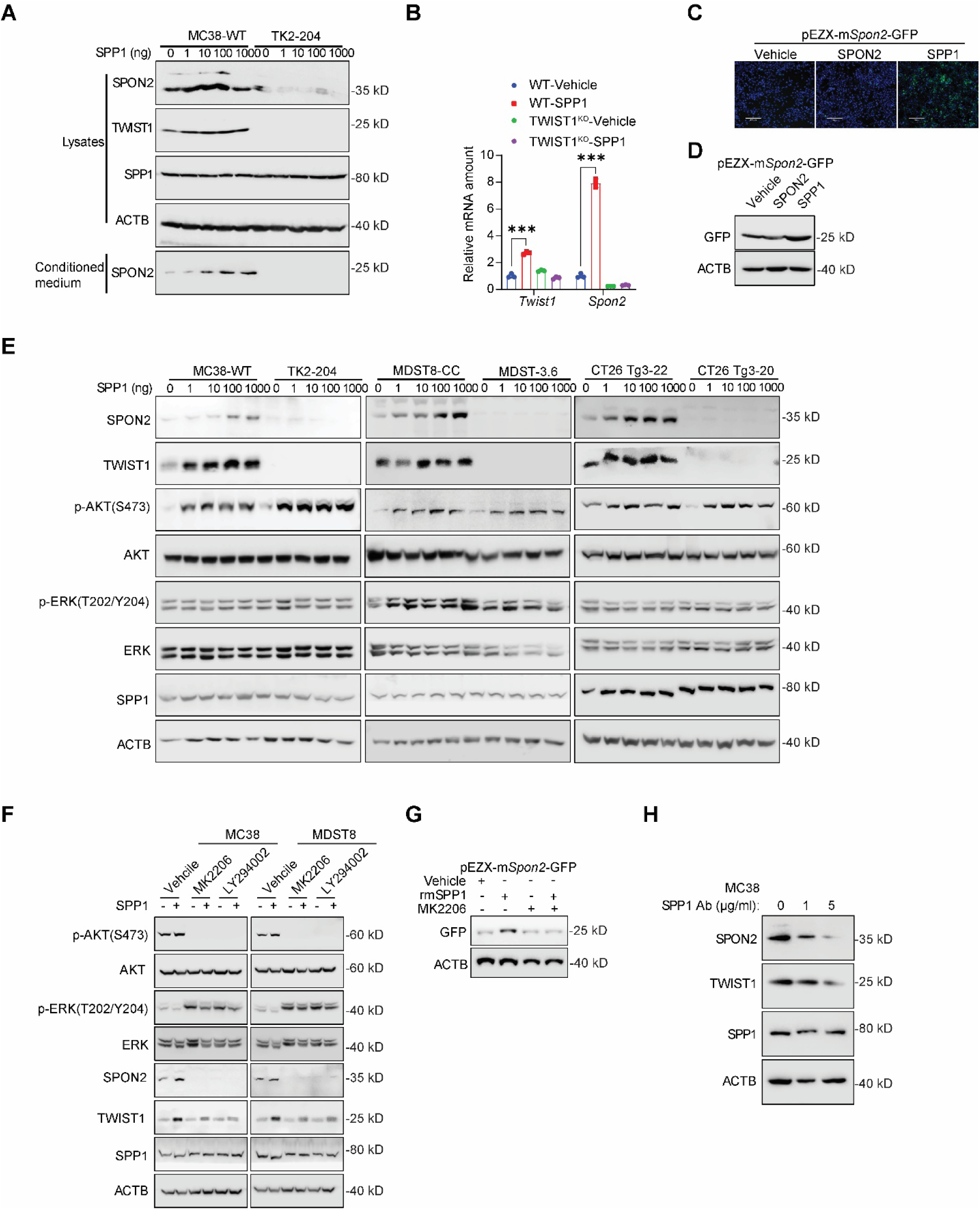
SPP1 enhances the TWIST1-SPON2 cascade. (A) Expression of TWIST1 and SPON2 in whole-cell lysates and SPON2 in the conditioned medium of MC38 cells following a pulse-chase with SPP1 protein for 24 hours at the indicated concentrations. MC38 WT and TWIST1 knockout cells (TK2-204) were starved for 48 hours prior to the SPP1 pulse-chase. (B) Expression of TWIST1 and SPON2 in MC38 cells treated with 100 ng/ml SPP1 protein for 24 hours. (C, D) Expression of *Spon2* promoter-driven GFP in MC38 cells treated with 100 ng/ml SPP1 protein for 24 hours. Representative GFP images are shown in (C), and Western blot analysis of protein expression is presented in (D). (E) Expression of TWIST1, p-AKT, p-ERK, SPP1, and SPON2 in whole-cell lysates following a pulse-chase with SPP1 protein for 24 hours at the indicated concentrations. MC38, MDST8, and CT26 cells with TWIST1 knockout were starved for 48 hours prior to the SPP1 pulse-chase. (F) Expression of TWIST1, p-AKT, p-ERK, SPP1, and SPON2 in whole-cell lysates following treatment with 100 ng/ml SPP1 protein and 1 µM PI3K/AKT inhibitors (MK2206 or LY294002) for 24 hours. (G) Expression of SPON2 promoter-driven GFP in MC38 cells treated with 100 ng/ml SPP1 protein and 1 µM MK2206 for 24 hours. (H) Expression of TWIST1 and SPON2 in MC38 cells treated with 1 or 5 µg/ml SPP1 monoclonal antibody for 24 hours. Results are expressed as mean ± SD. *P < 0.05, **P < 0.01, ***P < 0.001, and ****P < 0.0001.

Prior studies have reported that SPP1 regulates TWIST1 function by mediating signaling through the AKT and ERK pathways in breast cancer-associated fibroblasts [18]. In CRC cells, we demonstrated that SPP1 stimulation resulted in increased AKT phosphorylation without affecting ERK phosphorylation (Fig. 4E). Furthermore, inhibition of the PI3K/AKT pathway utilizing small molecule inhibitors, MK2206 and LY294002, reduced TWIST1 protein levels, along with concurrent decreases in *SPON2* mRNA and protein expression (Fig. 4F, S6B). The SPON2 promoter assay confirmed that AKT inhibition with MK2206 blocked SPP1-mediated *SPON2* transcription (Fig. 4G, S6C). Additionally, neutralization of secreted SPP1 with a specific antibody suppressed the expression of both TWIST1 and SPON2 proteins (Fig. 4H). Collectively, these results demonstrate that the TWIST1-SPON2 regulatory cascade is mediated through SPP1 in a PI3K/AKT-dependent manner.

The importance of SPON2 in metastatic CRC has been previously established, with various receptors, including α5β1, proposed to mediate the downstream effects of SPON2 [17]. Given that SPON2 plays a role in extracellular matrix organization and maintenance, we sought to investigate whether SPON2 secretion could stimulate the tumor microenvironment, including CAFs, within the CRC PM. Specifically, we investigated whether TWIST1 regulates the autocrine SPP1 signaling within cancer cells. Notably, comparison of wildtype cells to *TWIST1^KO^* cells revealed a significant decrease in secreted SPP1 protein levels, despite no alteration in cytoplasmic levels (Fig. 2J). Treatment of MC38 cells with recombinant Spon2 protein resulted in a significant increase in secreted SPP1 (Fig. 5A). Conversely, *TWIST1^KO^* cells maintained a modest level of suppressed secretion, even in setting of Spon2 stimulation (Fig. 5A), suggesting that the tumor cell-intrinsic SPP1 secretion mechanism is mediated downstream of TWIST1.

**Figure 5.**
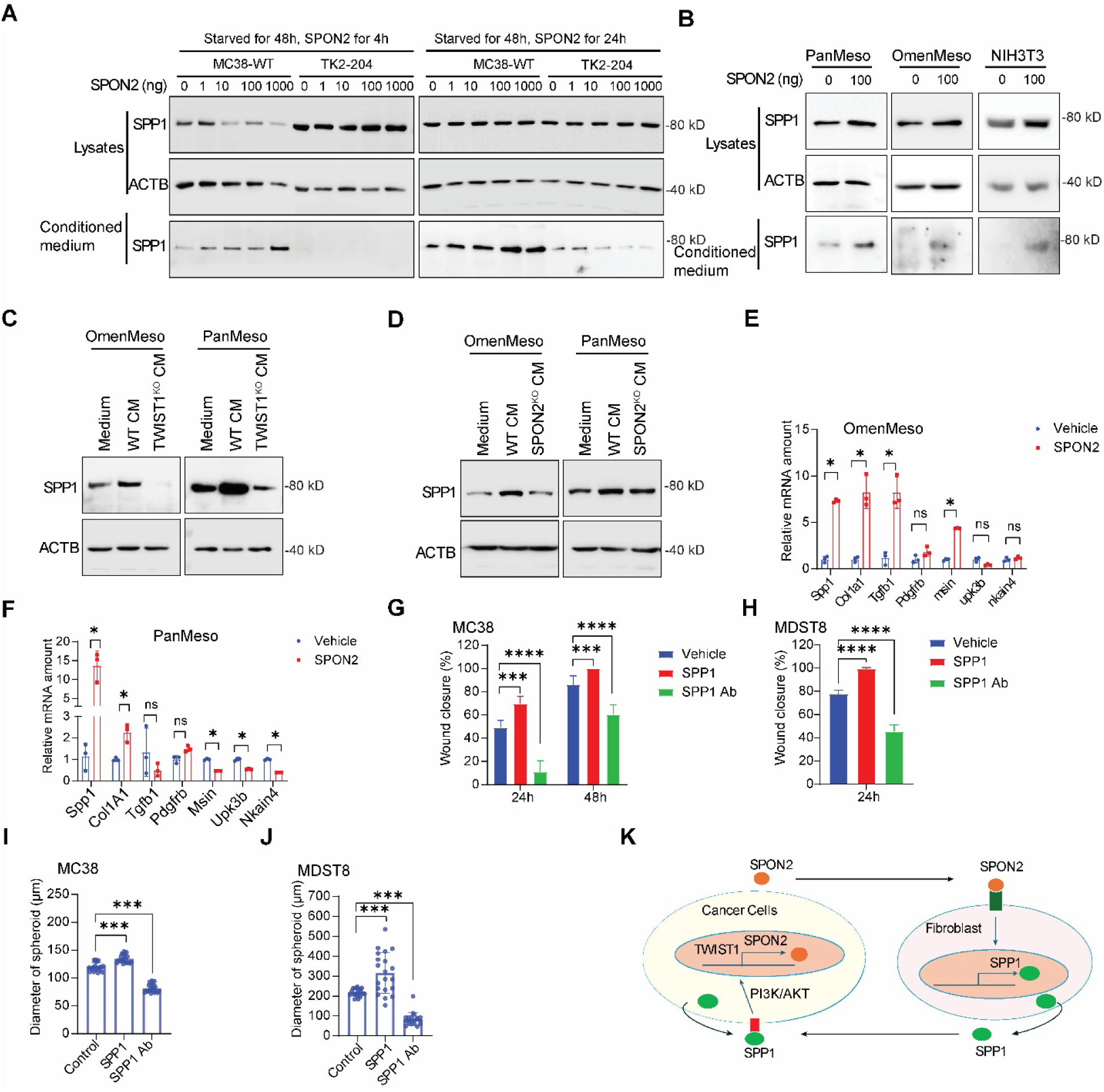
SPON2 regulates SPP1 expression and secretion to promote colon cancer cell migration, invasion, and stemness. (A) SPP1 expression in whole-cell lysates and conditioned medium of MC38 cells following a pulse-chase with SPON2 protein for 4 hours or 24 hours at the indicated concentrations. MC38 cells were starved for 48 hours prior to the SPON2 pulse-chase. (B) SPP1 expression in whole-cell lysates and conditioned medium of PanMeso, OmenMeso, and NIH-3T3 cells treated with 100 ng/ml SPON2 protein for 24 hours. (C) SPP1 expression in PanMeso and OmenMeso cells incubated with conditioned medium from MC38 TWIST1 knockout cells. (D) SPP1 expression in PanMeso and OmenMeso cells incubated with conditioned medium from MC38 SPON2 knockout cells. (E, F) Expression of mesothelial cell markers (Msin, Pdpn, Upk3b) and CAF markers (Spp1, Cola1, Tgti1, Pdgfrb) in OmenMeso (E) and PanMeso (F) cells treated with 100 ng/ml SPON2 protein for 24 hours. (G, H) Wound-healing assays to evaluate the migration of MC38 (G) and MDST8 (H) cells. MC38 and MDST8 cells were plated on Matrigel-coated plates with or without 100 ng/ml SPP1 protein or 1 µg/ml SPP1 monoclonal antibody. (I, J) Self-renewal capacity of MC38 (I) and MDST8 (J) cells cultured with 100 ng/ml SPP1 protein or 1 µg/ml SPP1 monoclonal antibody. (K) Schematic representation of the SPON2-SPP1 feedback loop. Results are expressed as mean ± SD. *P < 0.05, **P < 0.01, ***P < 0.001, and ***P < 0.0001.

Next, we examined whether tumor-derived secreted SPON2 influences stromal cells, particularly in regulating SPP1 secretion. Our recent study identified mesothelial cells—lining the peritoneal surface—as precursors to antigen-presenting CAFs (apCAFs) and serve as a major source of stromal SPP1 in CRC PM [9]. Given the critical role of tumor-mesothelial interactions in CRC PM progression, and the prominence of apCAFs within the CAF population, we investigated the effect of SPON2 on mesothelial cells [8]. We treated two prorietary mesothelial cell lines, PanMeso and OmenMeso—derived from pancreatic and omental mesothelium, respectively—with recombinant SPON2 or cancer cell-conditioned medium (CM). Treatment with SPON2 or CM from wildtype MC38 cells significantly increased SPP1 levels in mesothelial cells compared to treatment with CM from Twist*1^KO^* cells (Fig. 5B-C). Additionally, CM from Spon*2^KO^* cells resulted in reduced *Spp1* expression, suggesting SPON2 as an upstream regulator of *Spp1* in stromal cells (Fig. 5D). We then transfected OmenMeso cells with a *Spp1* promoter-driven *mCherry* reporter (pEZX-mSpp1-mCherry) and stimulated them with SPON2 protein. This stimulation led to enhanced mCherry expression, validating that SPON2 directly regulates *Spp1* transcription in stromal cells (Fig. S7A-C).

To further investigate whether tumor-secreted SPON2 mediates apCAF differentiation, we performed quantitative PCR analysis on a panel of CAF differentiation markers in PanMeso and OmenMeso cells following treatment with recombinant mSPON2. The results demonstrated that SPON2 significantly promoted the differentiation of PanMeso and OmenMeso into apCAFs, as evidenced by the upregulation of key CAF markers, including *Col1a1*, *Tgfb1*, *Pdgfrb*, and *Spp1* (Fig. 5E–F). This novel finding highlights that tumor-derived SPON2 not only induces SPP1 secretion in mesothelial cells within the tumor microenvironment but also plays a critical role in driving their differentiation into apCAFs.

To assess the functional consequences of SPON2-induced SPP1 expression, we treated wild-type MC38 and MDST8 cells with recombinant SPP1 protein and observed a significant increase in wound closure, migration, invasion, and sphere formation (Fig. 5G–J and S7D–G). Notably, neutralization of SPP1 using monoclonal antibodies markedly reduced these pro-tumorigenic properties, demonstrating that SPP1 is a key mediator of tumor cell aggressiveness (Fig. 5G–J and S7D–G). These findings further support the role of the TWIST1-SPON2 axis in driving SPP1 expression and secretion in stromal cells, thereby establishing a pro-tumorigenic microenvironment that facilitates colorectal cancer metastasis. Collectively, our results highlight SPON2 as a pivotal regulator of stromal remodeling and tumor progression, with potential implications for therapeutic targeting of the TWIST1-SPON2-SPP1 signaling axis in metastatic colorectal cancer.

Furthermore, SPP1-induced upregulation of *Spon2* was found to be TWIST1-dependent, indicating the presence of a positive feedback loop that reinforces TWIST1-SPON2 signaling and sustains a pro-metastatic tumor microenvironment in colorectal cancer (Fig. 4A, B). This suggests that the TWIST1-SPON2 axis actively drives *SPP1* expression and secretion in stromal cells, leading to an accumulation of SPP1 within the tumor microenvironment. In turn, this enriched SPP1 pool enhances PI3K/AKT-driven TWIST1-SPON2 signaling in cancer cells, ultimately promoting cancer cell invasion, enrichment of stem-like properties, and metastasis (Fig. 5K).

Finally, we established a syngeneic intraperitoneal peritoneal metastasis model using wild-type (WT) and *Spp1* knockout (*Spp1KO*) transgenic mice injected intraperitoneally with 50,000 MC38-WT or MC38-*Twist1KO* cells to validate the previous *in vitro* mechanistic insight *in vivo*. After 28 days, we assessed peritoneal tumor burden using the peritoneal carcinomatosis index (PCI) and evaluated ascites formation. In WT mice injected with MC38-WT cells, the highest tumor burden was observed, as reflected by significantly elevated PCI scores (Fig. 6A, B, and S8A). Additionally, these mice exhibited significant ascites accumulation (Fig. 6C). Interestingly, depletion of either stromal SPP1 or tumor-intrinsic TWIST1 led to a significant reduction in *Spp1*, *Spon2*, and *Col1a1* expression within the tumor, while concurrently increasing *Cd45* expression (Fig. 6D and S8B–E). These findings suggest that the SPP1-TWIST1-SPON2 signaling cascade contributes to cancer-associated fibrosis and immune evasion. Additionally, we assessed tumor burden using PCI scores and ascites volume in mice injected with 50,000 MC38-WT or MC38-*Spon2KO* cells to further investigate the impact of SPON2 on tumor progression. Loss of *Spon2* in MC38 cells resulted in a significant reduction in PCI scores, ascites formation, and cancer-associated fibrosis (Fig. 6E–H and S8F–G). Conversely, SPON2 depletion led to an increase in CD45-positive immune cell infiltration, suggesting that SPON2 contributes to the establishment of a fibrotic and immune cold tumor microenvironment (Fig. 6H). Collectively, these findings demonstrate that tumor-intrinsic TWIST1-SPON2 signaling is a key driver of colorectal cancer peritoneal metastasis by promoting stroma-derived SPP1 secretion. This axis fosters a pro-tumorigenic microenvironment characterized by enhanced fibrosis and reduced inflammation, highlighting its potential as a therapeutic target in metastatic colorectal cancer.

**Figure 6.**
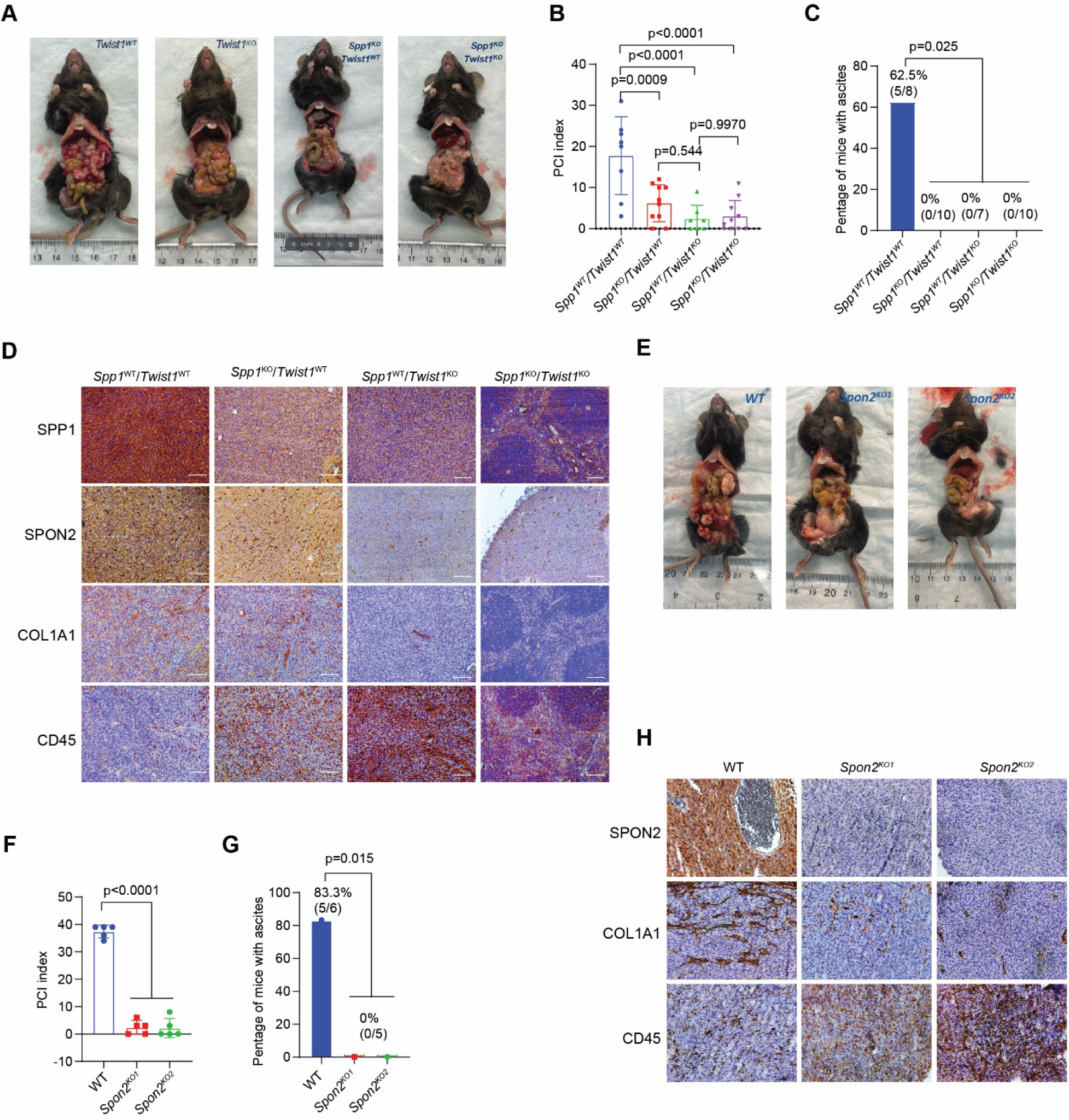
The TWIST1-SPON2-SPP1 cascade regulates peritoneal metastasis. (A) Representative images of peritoneal metastases in mice following intraperitoneal injection of MC38 cells. C57BL/6J mice, with or without *Spp1* gene knockout, were injected with 5 × 10⁴ MC38 cells, with or without *Twist1* knockout. Images were taken 28 days atier tumor cell injection. (B) Peritoneal carcinomatosis index (PCI) of mice that underwent intraperitoneal injection of MC38 cells. (C) Ascites volume in mice that underwent intraperitoneal injection of MC38 cells. (D) Immunohistochemical staining of SPP1, SPON2, COL1A1, and CD45 in tissue sections of MC38 peritoneal metastases. Scale bar: 200 µm. (E) Representative images of peritoneal metastases in mice following intraperitoneal injection of MC38 cells with *Spon2* knockout. Images were taken 28 days atier tumor cell injection. (F) Peritoneal carcinomatosis index (PCI) of mice that underwent intraperitoneal injection of 5 × 10⁴ MC38 cells with or without *Spon2* gene knockout in C57BL/6J mice. (G) Ascites volume in mice that underwent intraperitoneal injection of MC38 cells with *Spon2* knockout. (H) Immunohistochemical staining of SPP1, SPON2, COL1A1, and CD45 in tissue sections of MC38 peritoneal metastases. Scale bar: 200 µm. Results are expressed as mean ± SD. *P < 0.05, **P < 0.01, ***P < 0.001, and ***P < 0.0001.

## Discussion

Despite the high global incidence of colorectal cancer peritoneal metastases (CRC PM), there remains a significant gap in understanding its molecular mechanisms. This lack of progress contributes to rapid disease progression, frequent resistance to standard systemic therapies, and poor patient prognosis due to limited therapeutic options. Previous molecular profiling of CRC PM demonstrated an enrichment of the CMS4 subtype, which is associated with aggressive tumor behavior and poor outcomes[5]. The present study expands on this unique observation through an extensive genetic and molecular characterization of CRC PM with a novel identification of the SPP1-TWIST1-SPON2 paracrine and transcriptional axis as a distinctive driver of CRC PM tumorigenesis, a potential novel biomarker and an innovative therapeutic target. Dissection of the axis allowed for the discovery of an original molecular crosstalk within the tumor microenvironment between the tumor epithelium and CAFs, an important cellular circuit in CRC PM.

Our examination of the previously published large RNA-Seq data from CRC PM patients and validation within our own patient cohorts revealed a specific upregulation of a canonical EMT transcription factor, *TWIST1*. Emerging evidence highlights the crucial role of TWIST1 in CRC progression, particularly through its downstream gene activation responsible for metastasis and therapy resistance [19–22]. The correlation between high TWIST1 expression and advanced tumor stage, lymph node metastasis, and poor survival rates underscores its potential as a prognostic biomarker [23–25]. Moreover, TWIST1 has been implicated in the regulation of cancer stem cell (CSC) properties, which contribute to tumor initiation, recurrence, and resistance to conventional therapies, particularly to chemotherapeutic agents such as 5-fluorouracil (5-FU) and oxaliplatin [26]. Given its multifaceted role in CRC progression and specific upregulation within the CRC PM, our goal was to identify novel downstream gene(s).

Downregulation of TWIST1 through CRISP-Cas9 knockout or shRNA mediated silencing resulted in consistent phenotype with depleted metastatic potential as represented by decreased migration, Matrigel invasion, and wound closure assays. Moreover, the importance of TWIST1 in maintenance of CD44^+^ undifferentiated cancer stem cells in primary CRC was previously demonstrated [7]. Similarly, silencing of the TWIST1 expression decreased sphere formation potential, suggesting a loss of self-renewal capacity necessary to maintain the undifferentiated state. Despite the field’s current lack of the lineage tracing capabilities from primary tumor to metastatic cancer in CRC, we hypothesize that TWIST1 activation in subset of primary CRC results in downstream gene transcription that activates the invasion capacity, self-renewal capacity and differentiation for establishment of distant metastasis including PM. Moreover, the activation of TWIST1 specifically in CRC PM further supports the previous finding of CMS4 signature enrichment [5].

Notwithstanding the importance of TWIST1 in CRC PM, a comprehensive analysis of downstream target genes in CRC was not previously performed. Our extensive bioinformatics *in silico* analysis identified *SPON2*, among other dysregulated genes, as a potential biologically relevant target. In CRC, high *SPON2* expression was significantly correlated with worse disease-free survival and with advanced cancer stages. *SPON2* encodes a secreted protein with relatively unknown cell intrinsic and extrinsic functions. The assessment of *SPON2* in CRC PM and CRLM revealed a significant expression specificity mirroring *TWIST1* expression. Moreover, *SPON2^KO^* cells phenocopied *TWIST1^KO^* cells in assessment of intrinsic metastatic potentials via migration, Matrigel invasion, wound closure, and sphere formation assays. Analogous phenotype was also demonstrated by the antibody-mediated neutralization of SPON2 in cancer cells. Conversely, the addition of recombinant SPON2 or overexpressed SPON2 resulted in significant rescue of these deficient phenotypes. As such, the findings not only establish *SPON2* as a direct downstream TWIST1 target responsible for the metastatic phenotype in CRC PM tumorigenesis but also as a potential novel biomarker and therapeutic target. The cellular and molecular composition of TME serves to foster the malignant potential of specific cells to gain metastatic potential and exacerbate therapeutic resistance [27–29]. Peritoneum is a tissue that consists of various stromal cells, including mesothelial cells, fibroblasts, endothelial cells and immune cells. Our recent study has shown that mesothelial cells can differentiate into a specific CAF population, known as apCAF, to contribute to cancer progression [9]. CAFs can interact with tumor epithelium through paracrine signaling, such as TGF-β, to support aggressive cancer phenotypes, including CRC [7]. More importantly, CAF-derived SPP1 was recently identified to critically regulates the TME and maintain stemness of the cancer cells through CD44 [30]. Our investigation established that the mesothelial cells serve as an important source of stromal SPP1 in CRC PM. With previous establishment of CD44 as a receptor for SPP1 and of CD44 cells being enriched in TWIST1 expression, we sought to test the effect of SPP1 stimulation on downstream signaling and subsequent TWIST1 activation. The stimulation of tumor cells with SPP1 resulted in remarkable enhancement of metastatic potential. The primary activation of PI3K/AKT signaling cascade in our system revealed subsequent induction and amplification of TWIST1-*SPON2* transcriptional axis in tumor cells.

As a secreted factor, SPON2 likely exerts its biologic function through paracrine-mediated regulation of not only tumor cells but also within the TME. Stimulation with recombinant SPON2 or conditioned media containing SPON2 demonstrated increased SPP1 expression and secretion. More importantly, SPON2 stimulation initiated the mesothelial differentiation to apCAFs, thus demonstrating a novel paracrine signaling circuit between tumor cells and CAFs driving for CRC PM tumorigenesis. Our *in vivo* experiment further establishes this important novel circuit in CRC PM tumorigenesis and subsequent ascites production. Ascites production is frequently observed in PM and is a significant contributor of poor patient outcome due to abdominal pain, anorexia, nausea, and cachexia [31]. Commonly accepted hypothesis is that the tumor cells were primarily responsible for ascites production. Through our transgenic experiment, the stromal SPP1 secretion and subsequent SPON2 expression are likely responsible for the production of ascites. Following complete abolition of ascites in stromal *Spp1* knockout mice, we hypothesize that CAFs are the main activator of ascites production rather than the tumor epithelium.

In conclusion, our findings identify a novel SPP1-TWIST1-SPON2 axis that plays a critical role in the progression of CRC PM. This axis functions as both a key regulatory mechanism and a driver of tumor invasion and metastasis, fibrosis, and immune modulation within the peritoneal tumor microenvironment. Given its pivotal role, the SPP1-TWIST1-SPON2 axis holds significant potential as a biomarker for disease progression, an effector of metastatic signaling, and a promising therapeutic target for CRC PM intervention.

## Declarations

### Ethics approval and consent to participate

Eligible patients underwent informed consent at the Ohio State University Wexner Medical Center under IRB 2019C0139 and 2019C0196. Excess, deidentified tumor specimen were processed and entered a secure biorepository.

### Consent for publication

Not applicable

### Availability of data and materials

The datasets (GSE183202 and GSE50760) analyzed during the current study are available in the Gene Expression Omnibus under its identifier.

All data generated or analyzed during this study are available from the corresponding author on reasonable request.

### Author’s contributions

ZZ performed majority of the *in vitro* and *in vivo* studies. ALF performed majority of the bioinformatics analysis including ChIP-Seq and RNA-Seq. MHP generated shRNA knockdown and CRISPR-Cas9 knockout TWIST1 cell lines. XC assisted with the bioinformatics analysis. OE, AE, PMP, and JB assisted with the data analysis and manuscript writing. HH and ACK were responsible for hypothesis generation, experimental design, data analysis and manuscript writing.

## Acknowledgements

Not applicable

## Abbreviations

apCAF: Antigen presenting cancer associated fibroblast
CAF: Cancer-associated fibroblast
ChIP-Seq: Chromatin-immunoprecipitation-DNA sequencing
CM: Conditioned medium
CMS: Consensus molecular subtyping
CRC: Colorectal cancer
CRLM: Colorectal liver metastasis
EMT: Epithelial-mesenchymal transition
OmenMeso: Omental mesothelial cell
PanMeso: Pancreatic mesothelial cell
PM: Peritoneal metastasis

