## Supplemental Figures for "*TWIST1* mediated transcriptional activation of *SPON2* drives colorectal peritoneal metastasis through activation of cancer-associated fibroblast signaling network"

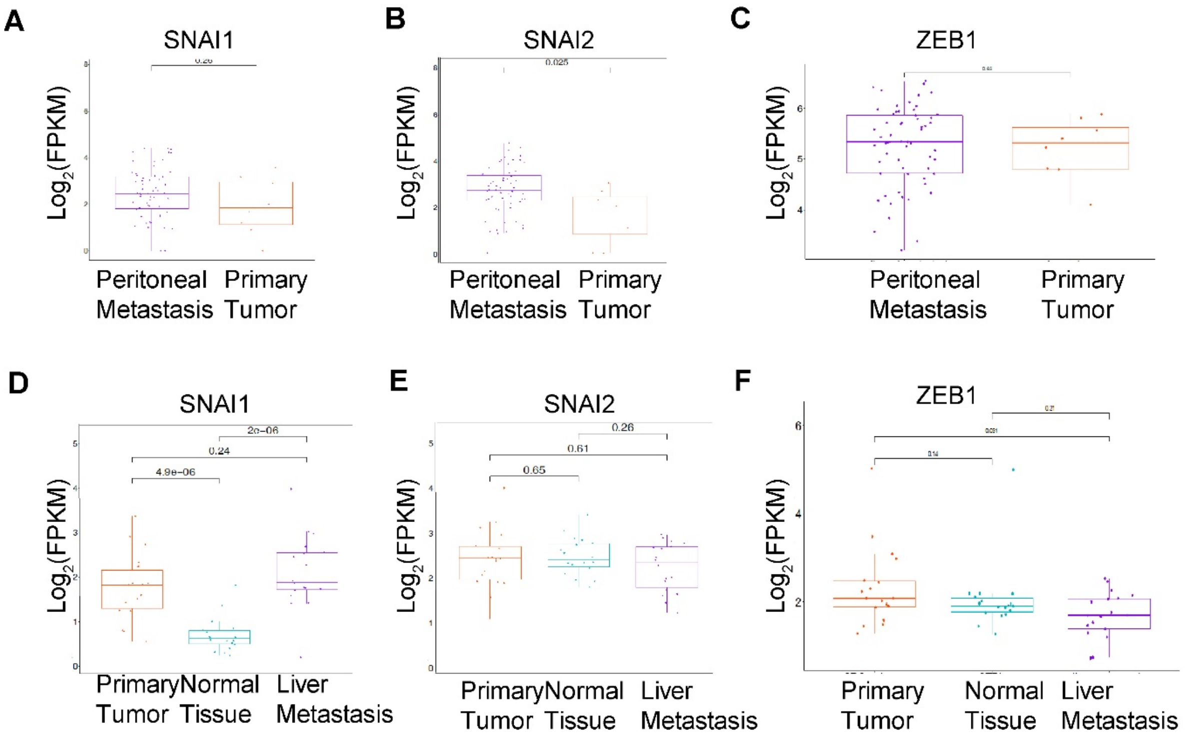


**Supplemental Figure 1.** Expression of EMT markers SNAI1, SNAI2, and ZEB1 in CRC peritoneal metastases (PM).

(A-C) Differential expression profiling of EMT markers (SNAI1, SNAI2, and ZEB1) in CRC peritoneal metastasis (PM) patient samples.

(D-E) Differential expression of EMT markers (SNAI1, SNAI2, and ZEB1) in CRC liver metastasis (LM) patient samples.

Results are expressed as mean ± SD. *P < 0.05, **P < 0.01, ***P < 0.001, and ***P < 0.0001.

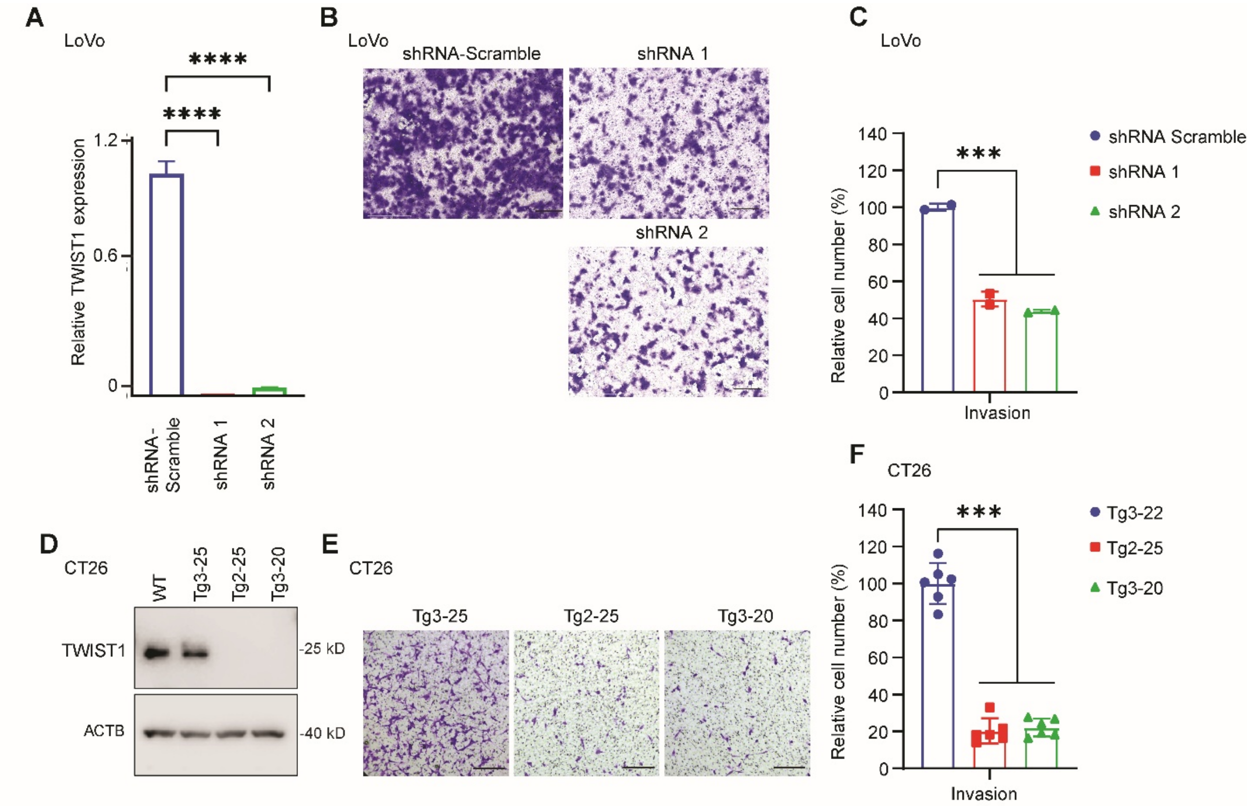


**Supplemental Figure 2.** TWIST1 regulates colon cancer cell migration, invasion, and stemness. (A) Expression of TWIST1 in LoVo cells with TWIST1 shRNA knockdown, assessed by western blot. (B-C) Transwell migration assays of LoVo cells with TWIST1 knockdown. Representative images are shown in (B), and quantification is presented in (C). Scale bar: 200 µm. (D) Expression of TWIST1 protein in CT26 TWIST1 knockout single colony clones, confirmed by western blot. (E-F) Transwell migration assays of CT26 cells with TWIST1 knockout. Representative images are shown in (E), and quantification is presented in (F). Scale bar: 200 µm. Results are expressed as mean ± SD. *P < 0.05, **P < 0.01, ***P < 0.001, and ***P < 0.0001.

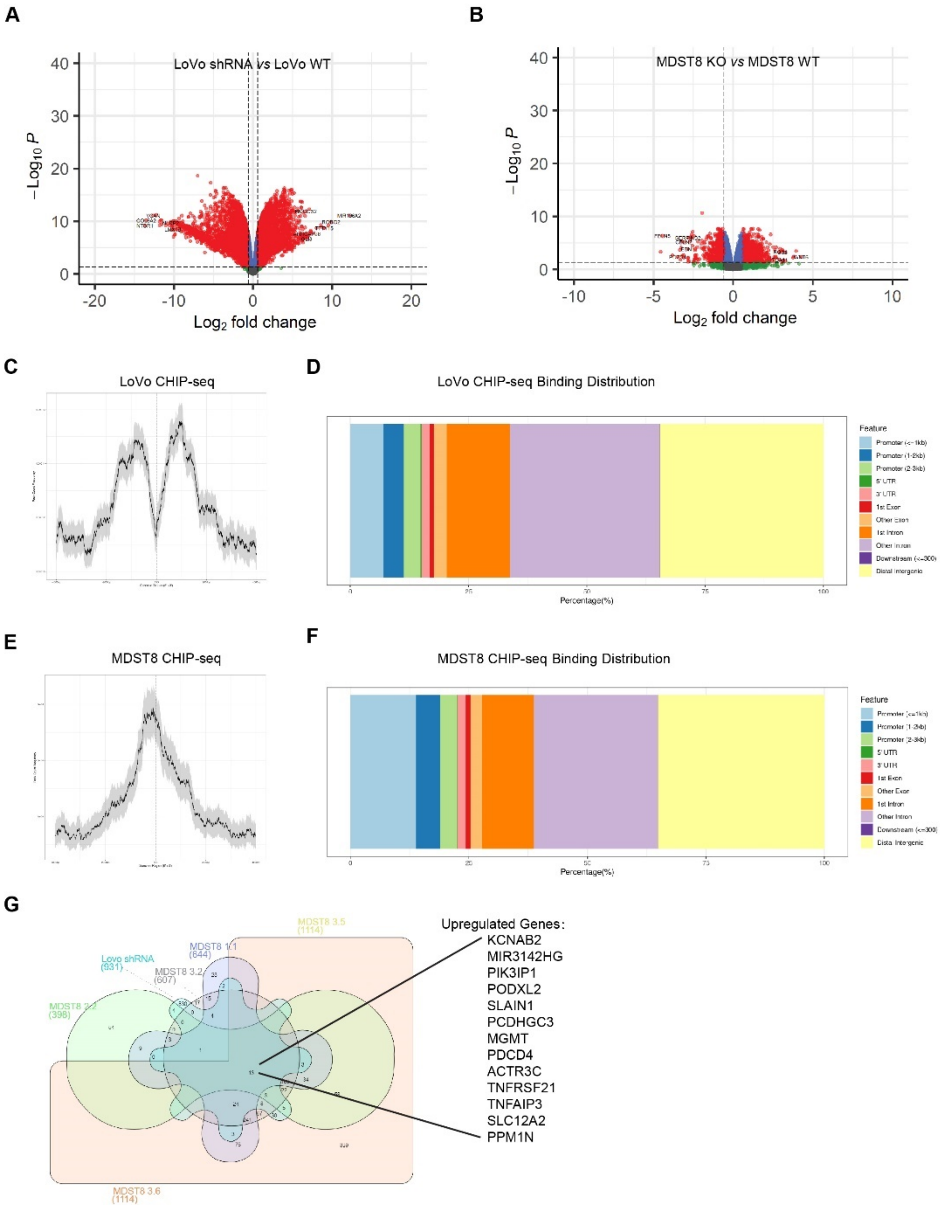


**Supplemental Figure 3.** RNA-seq and TWIST1 ChIP-seq analysis of LoVo and MDST8 cells.

(A) Volcano plot depicting differentially expressed genes in LoVo cells with TWIST1 knockdown (shRNA-1) versus LoVo wild-type (WT) cells.

(B) Volcano plot depicting differentially expressed genes in MDST8 TWIST1 knockout (clone 3.6) versus MDST8 WT cells.

(C) Peak count distribution of TWIST1 binding relative to the genome in LoVo cells, based on ChIP-seq analysis.

(D) Genomic feature distribution of TWIST1 binding sites in LoVo cells.

(E) Peak count distribution of TWIST1 binding relative to the genome in MDST8 cells, based on ChIP-seq analysis.

(F) Genomic feature distribution of TWIST1 binding sites in MDST8 cells.

(G) Overlay of ChIP-seq and RNA-seq data from LoVo and MDST8 cells. Venn diagrams illustrate genes with TWIST1-bound promoters (from ChIP-seq) that are upregulated in TWIST1-deficient cells, highlighting key TWIST1-regulated targets.
Results provide insight into the transcriptional and epigenetic regulation mediated by TWIST1 in colorectal cancer metastasis.

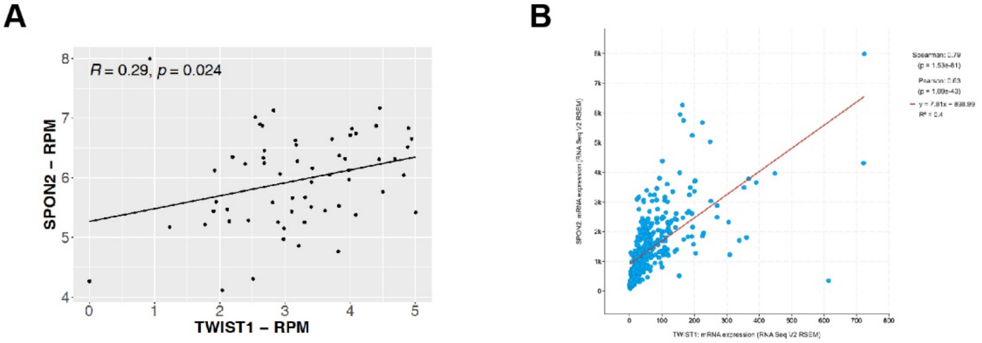


**C**

| SPON2 in COAD (n=458) | | | | | | |
|---|---|---|---|---|---|---|
| Stage | coef | HR | se(coef) | 95%CI | z | p |
| 1 | 0.004 | 1.004 | 0.093 | 0.837-1.206 | 0.047 | 0.962 |
| 2 | 0.754 | 2.125 | 0.54 | 0.737-6.124 | 1.396 | 0.163 |
| 3 | 1.312 | 3.712 | 0.537 | 1.296-10.636 | 2.443 | 0.015 |
| 4 | 2.369 | 10.692 | 0.535 | 3.746-30.512 | 4.429 | 0.000 |

$R^2$=0.1
Likelihood ratio test p=4.29e-0.9
Wald test p=1.63e-09
Score (logrank) test p=7.16e-12

**Supplemental Figure 4.** SPON2 Expression as a Poor Prognostic Marker in PM Patient Samples and Its Correlation with TWIST1 Expression

(A) Significant positive correlation between TWIST1 and SPON2 expression in CRC peritoneal metastasis (PM) patient samples, indicating a potential regulatory relationship.

(B) TCGA data analysis showing a significant positive correlation between TWIST1 and SPON2 expression in CRC patients, further supporting the association observed in PM samples.

(C) Increasing hazard ratio for TCGA colon cancer samples, stratified by stage, demonstrating that higher SPON2 expression is associated with worse prognosis across disease stages.

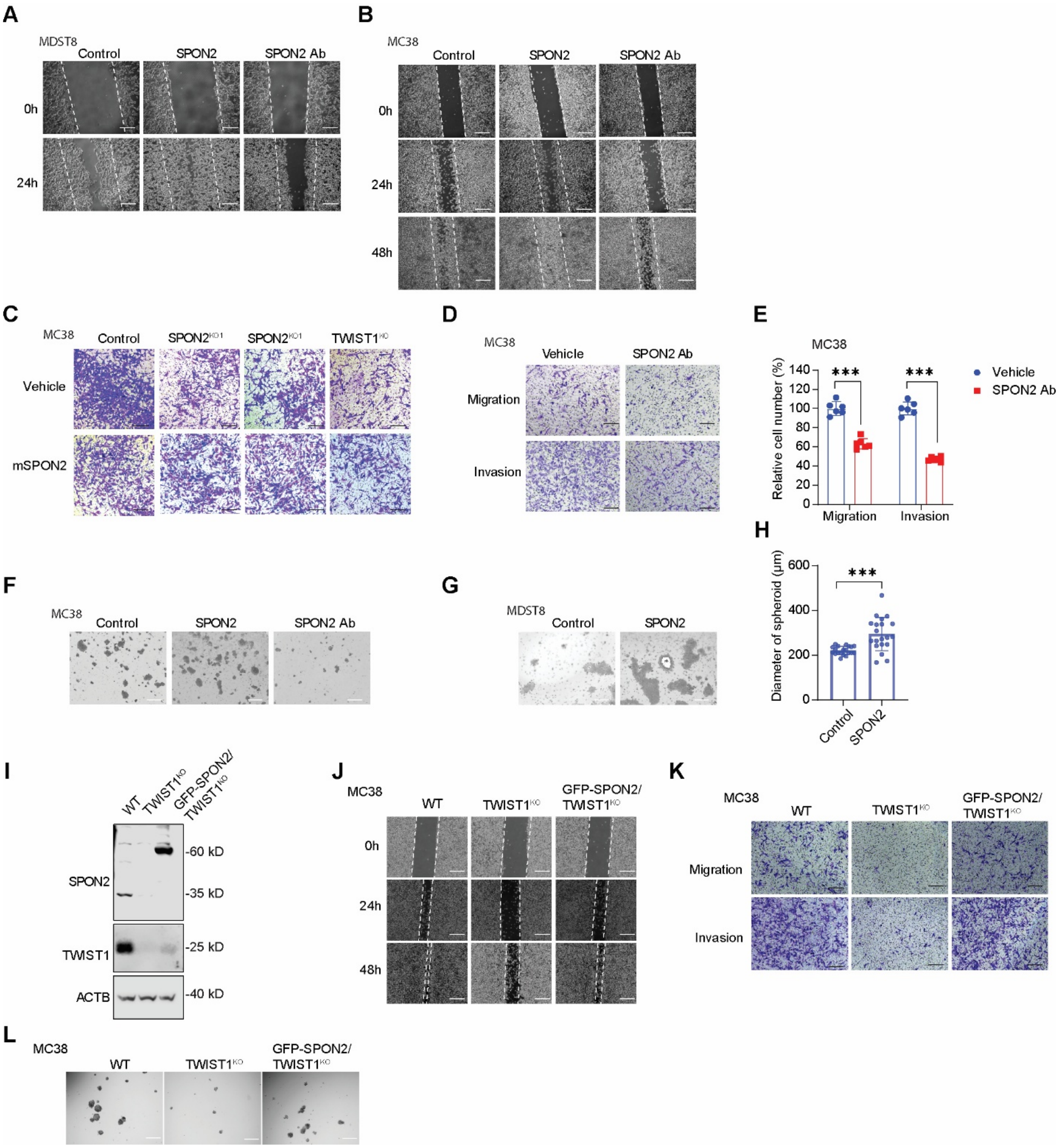


**Supplemental Figure 5.** The TWIST1-SPON2 Cascade Regulates Colon Cancer Cell Migration, Invasion, and Stemness

(A-B) Wound-healing assay to assess the migration of MC38 (A) and MDST8 (B) cells, plated on Matrigel-coated plates with or without 100 ng/ml SPON2 protein or 1 µg/ml SPON2 monoclonal antibody. Representative images shown. Scale bar, 200 µm.

(C) Transwell Matrigel invasion assays to evaluate the effect of SPON2 protein on the chemoinvasion of MC38 TWIST1 or SPON2 knockout cells. The Transwell was coated with 1 mg/ml Matrigel with or without 100 ng/ml SPON2 protein and incubated for 48 hours with 10% FBS. Representative images shown. Scale bar, 200 µm.

(D-E) Transwell migration and Matrigel invasion assays to assess haptotactic migration and Matrigel chemoinvasion in MC38 cells with or without 100 ng/ml SPON2 protein. Representative images are shown in (D), and statistical results are presented in (E). Scale bar, 200 µm.
(F) Assessment of self-renewal capacity in MC38 cells cultured with 100 ng/ml SPON2 protein or 1 µg/ml SPON2 monoclonal antibody.
(G-H) Assessment of self-renewal capacity in MDST8 cells cultured with 100 ng/ml SPON2 protein. Representative images shown in (G) and statistical results shown in (H). Scale bar, 200 µm.
(I) Western blot showing the expression of GFP-SPON2 in MC38 cells with TWIST1 knockout.
(J) Wound-healing assay to assess the migration of MC38 TWIST1 knockout cells with overexpression of GFP-SPON2. MC38 cells were plated on Matrigel-coated plates.
(K) Transwell migration and Matrigel invasion assays to assess haptotactic migration and Matrigel chemoinvasion of MC38 TWIST1 knockout cells with overexpression of GFP-SPON2.
(L) Self-renewal capacity of MC38 TWIST1 knockout cells with overexpression of GFP-SPON2.
Results are expressed as mean ± SD. *$P < 0.05$, **$P < 0.01$, ***$P < 0.001$, and ****$P < 0.0001$.

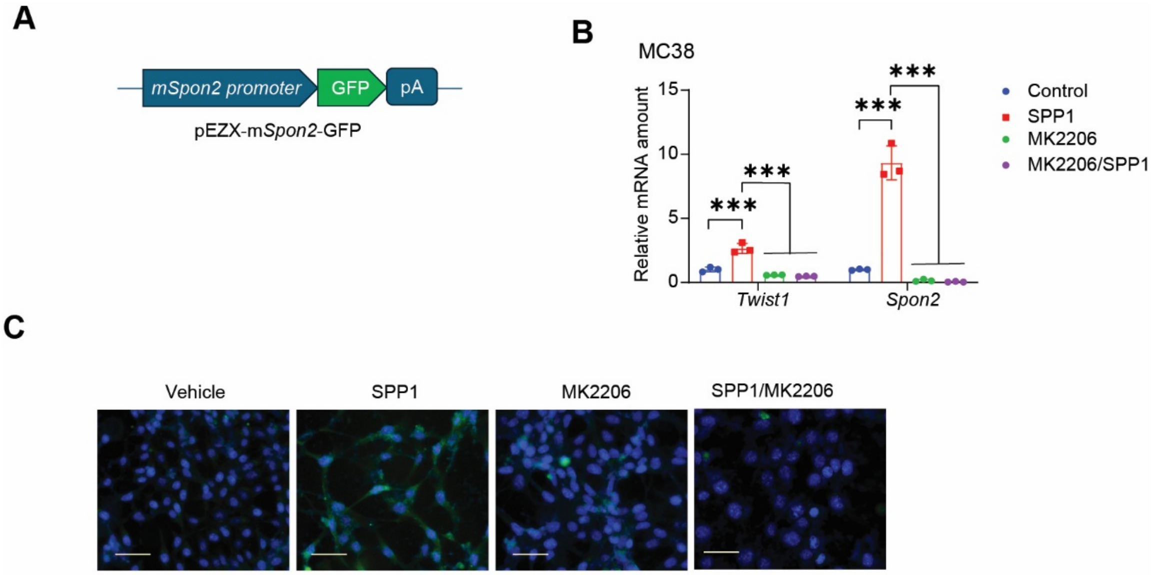


**Supplemental Figure 6.** The SPP1 Enhances TWIST1-SPON2 Cascades

(A) The schematic figure shows the structure of p-EZX-m*Spon2*-GFP.

(B) Expression of Twist1 and Spon2 mRNA in MC38 cells treated with 100 ng/ml SPP1 protein with or without 1 µM MK2206 for 24 hours.

(C) Expression of Spon2 promoter-GFP in MC38 cells treated with 100 ng/ml SPP1 protein and 1 µM MK2206 for 24 hours.

Results are expressed as mean ± SD. Statistical significance was determined with *P < 0.05, **P < 0.01, ***P < 0.001, and ****P < 0.0001.

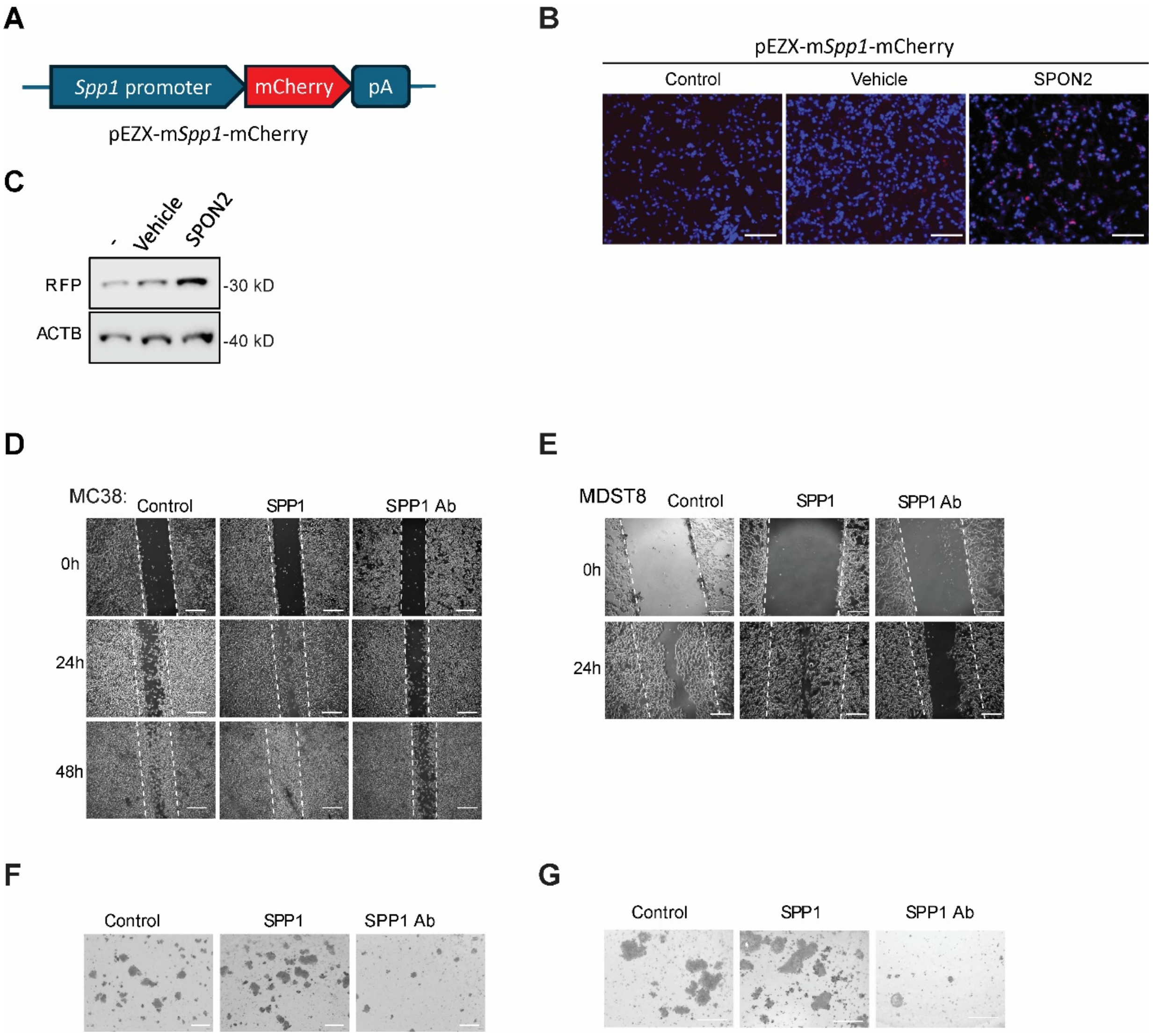


**Supplemental Figure 7.** The SPON2 increases SPP1 Expression in Mesothelial Cells and Enhances Colon Cancer Cell Migration, Invasion, and Stemness

(A) The schematic figure shows the structure of p-EZX-m*Spp1*-mCherry.

(B-C) Expression of Spp1 promoter-mCherry in OmenMeso cells treated with 100 ng/ml SPON2 protein for 24 hours. Representative images are shown in (B), and Western blotting results are shown in (C).

(D-E) Wound-healing assays to evaluate the migration of MC38 (D) and MDST8 (E) cells. MC38 and MDST8 cells were plated on Matrigel-coated plates with or without 100 ng/ml SPP1 protein or 1 μg/ml SPP1 monoclonal antibody.

(F-G) Self-renewal capacity of MC38 (F) and MDST8 (G) cells cultured with 100 ng/ml SPP1 protein or 1 μg/ml SPP1 monoclonal antibody.

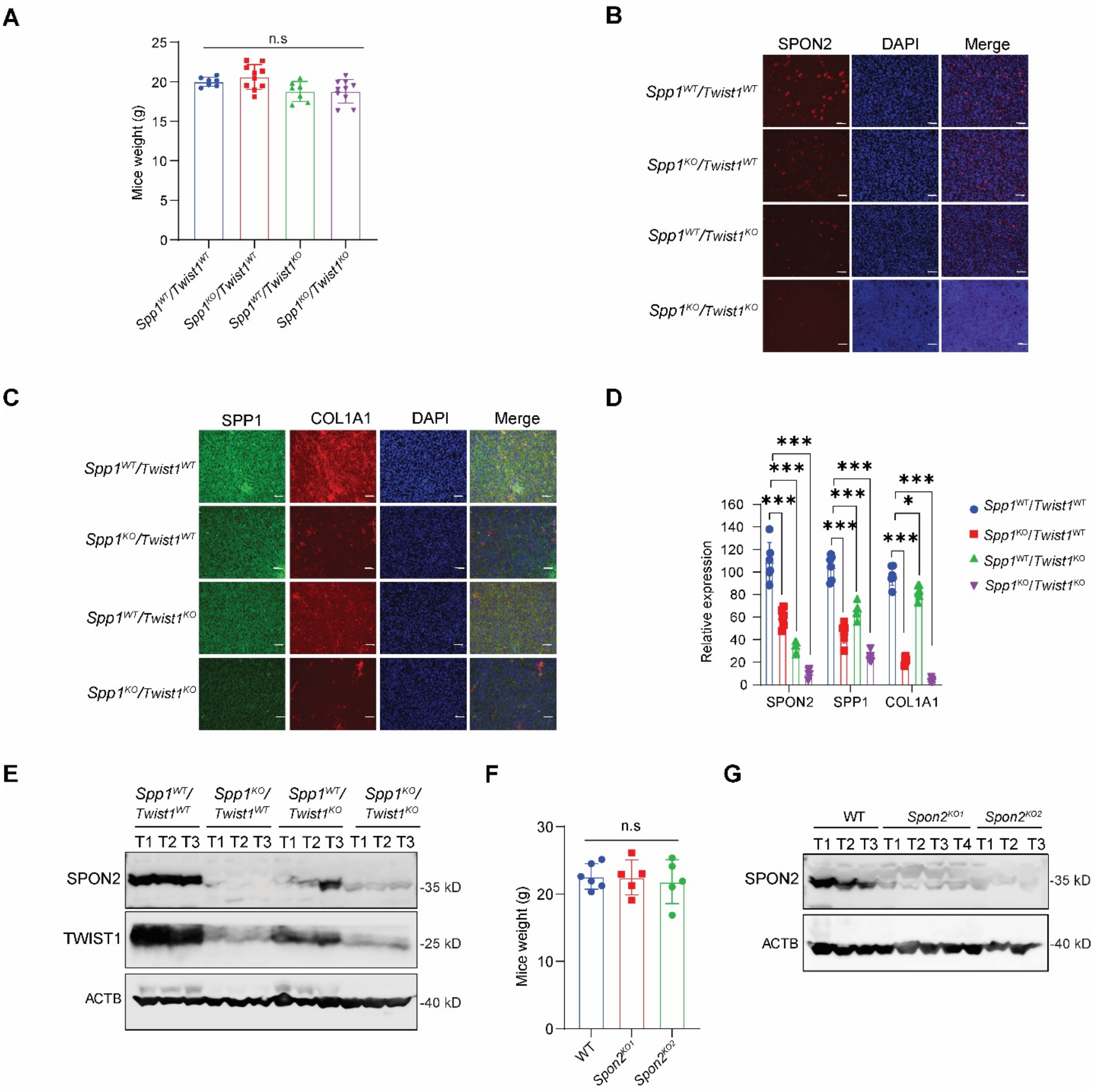


**Supplemental Figure 8.** The TWIST1-SPON2-SPP1 Cascades Regulate Peritoneal Metastasis

(A) Mouse weight 28 days after tumor cell injection. C57BL/6J mice, with or without *Spp1* gene knockout, were injected with 5 x 10^4 MC38 cells, with or without *Twist1* knockout.

(B) Immunofluorescence staining of SPON2 expression in tissue sections of MC38 peritoneal metastases.

(C) Immunofluorescence staining of SPP1 and COL1A1 expression in tissue sections of MC38 peritoneal metastases. Scale bar, 50 µm.

(D) Quantification of SPON2, SPP1, and COL1A1 expression was performed using immunofluorescence staining. Data was collected from six independent images per sample group.

(E) Western blotting of SPON2 and TWIST1 expression in MC38 peritoneal metastases in C57BL/6J mice, with or without *Spp1* gene knockout.

(F) Mouse weight 28 days after MC38 cells with or without *Spp1* gene knockout injection in C57BL/6J mice.
(G) Western blotting of SPON2 and TWIST1 expression in MC38 peritoneal metastases.
